# PARROT: Phase-Altering Regulatory Rewiring Over Time

**DOI:** 10.64898/2026.06.24.734262

**Authors:** Chen Chen, Megha Padi, John Quackenbush

**Affiliations:** Department of Biostatistics, Harvard T.H. Chan School of Public Health, USA; Department of Molecular and Cellular Biology, University of Arizona, USA

**Keywords:** change-point detection, dynamic networks, stochastic block model, gene regulatory networks, community detection

## Abstract

**Motivation:** Gene regulatory networks undergo dynamic restructuring during development and disease. Identifying when and how these networks change is crucial for understanding developmental and disease transitions, yet existing change-point detection methods often ignore network structure or lack interpretable community assignments.

**Results:** We present PARROT (Phase-Altering Regulatory Rewiring Over Time), a framework for detecting change-points in dynamic networks using Stochastic Block Models. PARROT jointly estimates change-point locations and community structure across four network classes: unipartite and bipartite with either Gaussian or Bernoulli edge models. Simulations demonstrate improved performance and community recovery compared to other methods. Applications to human cardiac differentiation and mouse lung development data successfully recovered known phase boundaries. PARROT identifies both which genes are reassigned across modules and how the connections change between states.

**Availability:** PARROT is available as an R package at https://github.com/cchen22/PARROT.

**Supplementary information:** Supplementary data are available at Bioinformatics online.

## 1. Introduction

Gene regulatory networks (GRNs) are inherently dynamic, undergoing restructuring during development, cellular differentiation, disease progression, and aging (Barabási and Oltvai, 2004; Karlebach and Shamir, 2008; Schlitt and Brazma, 2007).

Determining *when* these transitions occur and *which* regulatory programs are affected is fundamental to understanding dynamic biological processes. While numerous methods exist for detecting change-points (CPs) in time series data (Killick et al., 2012; Fryzlewicz, 2014; Truong et al., 2020) and for community detection in networks (Karrer and Newman, 2011; Padi and Quackenbush, 2018), few approaches address the joint problem of detecting regulatory transitions in dynamic network data that alter biological processes through network rewiring to change phenotypic state.

Commonly used network change-point detection methods fall into several categories. Graph-based methods like gSeg (Chen et al., 2020) use distribution-free scan statistics but do not recover community structure. Bayesian approaches such as NetworkChange (Park and Sohn, 2022) provide uncertainty quantification but are computationally intensive and limited to unipartite networks, whereas regulatory networks are typically bipartite, with transcription factor (TF) nodes and target gene nodes having very different properties. Energy-based methods (James and Matteson, 2015) require vectorization of network data, losing structural information. From the other direction, dynamic community detection methods such as PisCES (Liu et al., 2018) apply spectral clustering with eigenvector smoothing to recover *persistent* community structure across time, but flag change-points only implicitly, with no tested location, p-value, or bipartite network support. None of these jointly estimates change-point locations *and* community membership in a statistically robust framework, leaving researchers unable to identify *when* network rewire and *which* modules drive those transitions.

Here we present PARROT (Phase-Altering Regulatory Rewiring Over Time), a model-based framework that overcomes these limitations. PARROT uses Stochastic Block Models (SBMs) (Holland et al., 1983) to model network structure and uses a profile-likelihood scan and permutation testing to detect change-points. In PARROT, a change-point corresponds to a time at which the SBM parameters shift: the community memberships (which genes and TFs belong to each module), the block connectivity (how strongly modules interact), or both. Key innovations include: (1) support for weighted bipartite networks, enabling analysis of transcription factor (TF)-to-gene regulatory networks; (2) joint estimation of change-point locations and community assignments; (3) Wild Binary Segmentation (Fryzlewicz, 2014) for multiple change-point detection; and (4) C++-accelerated Variational Expectation–Maximization (VEM) for computational efficiency.

We validate PARROT through comprehensive simulations and apply it to two biological network series: (1) a TF– gene network derived from scRNA-seq data obtained from a human induced pluripotent stem cell (hiPSC) cardiac differentiation series, and (2) a co-expression network series obtained from a mouse postnatal lung development experiment. Both applications demonstrate PARROT’s ability to reveal discrete network transitions with mechanistic interpretability.

## 2. Methods

### 2.1 Model formulation and inference

Let **Y** = {*Y* ^(1)^, …, *Y* ^(*T*)^} denote an ordered sequence of *T* network snapshots, where each 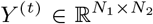 represents the adjacency matrix at time *t*. For unipartite networks, *N*_1_ = *N*_2_ = *N* ; for bipartite networks, such as TF→gene regulatory networks (Glass et al., 2013) or SNP→gene eQTL networks (Platig et al., 2016), *N*_1_ ≠*N*_2_.

#### Stochastic Block Model

Each node is assigned to one of *Q* communities. The edge probability/weight between nodes in communities *q* and *r* is governed by block parameters *γ*_*qr*_:

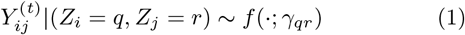

where *f* is Gaussian (for weighted networks) or Bernoulli (for binary networks), and *Z*_*i*_ denotes node *i*’s community assignment.

#### Change-Point Model

Under the alternative hypothesis of a change point at time *τ*, networks follow SBM parameters *θ*_1_ for *t* ≤ *τ* and *θ*_2_ for *t > τ*, where for each segment *k* ∈ {1, 2} the parameter bundle *θ*_*k*_ = (***π***_*k*_, {*γ*_*qr,k*_}) collects the mixing proportions ***π***_*k*_ and the block parameters *γ*_*qr,k*_ defined in Eq. (1) (*γ*_*qr,k*_ = *μ*_*qr,k*_ for Gaussian and *γ*_*qr,k*_ = *p*_*qr,k*_ for Bernoulli networks):

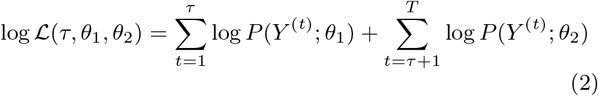

The change point is estimated as:

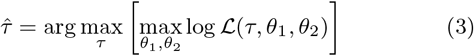

#### Model fitting and approximation

We fit SBMs using VEM and maximizing the Evidence Lower Bound (ELBO), which serves as a tractable surrogate for the intractable marginal log-likelihood (Supplementary Methods). For a candidate change-point *τ*, we evaluate the profile objective by summing ELBOs from the left and right segments and selecting the maximizer 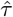. PARROT reports the split with the *largest* profile objective (best-supported partition), while local minima correspond to poorly supported splits. For temporally long sequences, PARROT also provides an optional score-based scan as a faster approximate alternative to full profile refitting (Supplementary Table S6; Supplementary Figure S3).

#### Profile likelihood confidence intervals

Let *ℓ*(*τ*) denote the profiled objective at split *τ* (left-segment ELBO plus right-segment ELBO, each maximized over segment-specific SBM parameters), and let 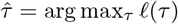. We form a likelihood-ratio-type confidence set by retaining indices that satisfy

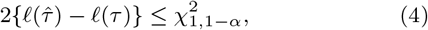

or equivalently 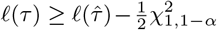. The degree of freedom is 1 because the scanned change-point location is one-dimensional. In our implementation, this is a large-sample chi-square cutoff approximation applied to a discrete split index and an ELBO surrogate (not the exact marginal log-likelihood), so that interval coverage should be interpreted as approximate in finite samples.

#### Likelihood Ratio Test and P-values

Define the likelihood ratio statistic as

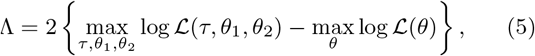

where the first term is the maximized log-likelihood under the alternative (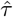) and the second is the null (no change). Because optimization is based on an ELBO surrogate rather than the exact marginal likelihood, we emphasize resampling-based p-values (permutation/bootstrap) for practical inference.

By default, we use a permutation test. We randomly permute the time labels of the observed networks to generate *B* null replicates 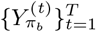, refit the change-point model on each, and compute

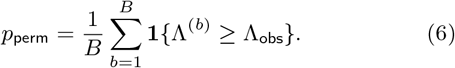

Parametric bootstrap p-values are also available: we simulate networks from the fitted null SBM and compare Λ_obs_ to the bootstrap distribution of Λ. Both permutation and bootstrap tests require additional refits and do not provide exact finite-sample guarantees under all temporal dependence settings.

### 2.2 Multiple change-point search

For detecting multiple change-points, we use Wild Binary Segmentation (WBS) (Fryzlewicz, 2014). In implementation, WBS generates candidate split points from random intervals using ELBO-gain scans, then applies priority-ordered filtering with minimum-segment separation and a maximum-number-of-change-points constraint. This candidate-filtering strategy is practical and reproducible, but it is not a formal global error-rate controlling stopping rule. Additionally, PARROT supports Binary Segmentation (BS) and PELT (Pruned Exact Linear Time) (Killick et al., 2012) methods as alternative multiple change-point search strategies. Two candidate-scanning modes are available. The *score scan* performs a single global (null) SBM fit, freezes variational memberships, and precomputes cumulative block sufficient statistics; each candidate split *τ* is then scored by differencing these cumulative sums, giving *O*(1) updates per position without per-split variational refits. This acceleration is in the spirit of CUSUM-style cumulative-sum scanning (Page, 1954; Truong et al., 2020). The *profile scan* instead evaluates the profiled ELBO *ℓ*(*τ*) by refitting separate SBMs on the left and right segments at every candidate split, typically improving fidelity at higher computational cost. Practical guidance on scan-mode choice, CI interpretation, p-value assumptions, and sample-size/runtime tradeoffs is summarized in Supplementary Table S6.

### 2.3 Software implementation

PARROT is implemented in R with C++ acceleration via Rcpp. The core VEM algorithm achieves *O*(*T NQ*^2^) complexity per iteration. The package supports all four network types, automatic *Q* selection using Integrated Classification Likelihood (ICL), and multiple inference methods.

### 2.4 Simulated network construction

We generated a sequence of SBM snapshots with a single change-point *τ* ^*\**^ separating two parameter regimes, considering all four network classes supported by PARROT: unipartite/bipartite × Gaussian/Bernoulli. Across every class we used *T* = 10 time points, *Q* = 2 communities, *τ* ^*\**^ = 5, and *n* = 20 replicates per configuration; node counts were *N* = 12 for unipartite and *N*_1_ × *N*_2_ = 8 × 10 for bipartite networks. These deliberately small/short settings (i) avoid ceiling effects so that competing methods retain non-trivial discrimination, and (ii) stress-test the ability of every method to recover a change with limited evidence per segment. Three transition families were simulated: (i) *global mean shift* (block parameters change uniformly while memberships are held fixed; relatively weak signal), (ii) *community swap* (block parameters held fixed while ∼35% of nodes switch community membership), and (iii) *mixed* (a moderate global shift combined with ∼30% community reassignment and rewiring). The exact block matrices, noise scales, shift magnitudes, and swap fractions are listed in Supplementary Table S1; the matching tolerance and other evaluation metrics are listed in Supplementary Table S2. To verify that PARROT’s underlying SBM fitting recovers parameters under static (no-CP) conditions on *larger* networks, we additionally ran *N* = 40 unipartite and 15 × 20 bipartite static simulations with *T* = 20 (Supplementary Figure S1).

### 2.5 Data preprocessing and network construction

The two real-data applications presented here analyze a series of networks derived from publicly available time course datasets archived in GEO (Edgar et al., 2002).

We used scRNA-seq data collected from a time course (GEO accession GSE202398) that profiled the WTC and SCVI111 hiPSC lines across 12 time points (days 0–7, 11, 13, 15, 30) that include a Day2-induced cardiac differentiation under a biphasic small-molecule WNT protocol (Galdos et al., 2023). Raw 10x gene expression matrices were demultiplexed using run-specific hashtag oligo metadata, per-sample cells were quality filtered based on feature count and mitochondrial fraction and these were then downsampled to balance snapshots. Using these data, we constructed an expanded WNT-focused bipartite feature set (25 TFs and 150 target genes) by selecting variance-ranked genes with cardiac-regulatory TF seeds, reducing sparsity and noise while preserving key developmental regulators. For each day, TF and target expression were standardized across cells, Spearman TF–target correlation edge weights were computed; these were transformed using the Fisher *z* transformation, *z* = atanh(*r*), and averaged across two cell lines to form one weighted bipartite network snapshot per day (*T* = 12). We analyzed this time-ordered network series using PARROT in single-CP mode. The protocol-defined reference CP is the WNT-activation to WNT-inhibition switch boundary (Day2→Day3), used as an external intervention-grounded reference.

As a second example, we chose a mouse postnatal lung development dataset (GEO accession GSE74243). We used processed RMA-normalized log2 microarray expression data from whole lung tissue across three strains, A/J, C57BL/6J, and C3H/HeJ (Beauchemin et al., 2016). We parsed strain and postnatal day from sample labels, mapped probes to gene symbols, and collapsed duplicates by mean expression. Building on the five postnatal molecular stages from Beauchemin et al. (2016)—ALV1 (P0–P3), ALV2 (P4–P5), ALV3 (P7–P13), ALV4 (P14–P18), and MAT (P21–P56)—we subdivided ALV3 and MAT to increase temporal resolution, yielding seven time points: T1=P0–P3, T2=P4–P5, T3=P7–P9, T4=P11– P13, T5=P14–P18, T6=P21–P24, T7=P30–P56. Reference boundaries include *T* = 5 (ALV4→MAT: onset of homeostatic mature expression) and potentially *T* = 2–4 (alveolarization plateau transitions). Importantly, these stages were defined via unsupervised PCA of expression levels (Beauchemin et al., 2016) and were thus independent of network inference and analysis; this PARROT-based analysis therefore tests whether co-expression *network structure* transitions align with expression-derived boundaries.

For each time point, we merged replicates across strains using ComBat (Johnson et al., 2007) and COBRA (Micheletti et al., 2024) to remove both first-order and second-order batch effects while preserving biological signal, thereby maximizing sample size, selected the top 100 most variable genes to construct the co-expression network, computed Spearman gene– gene correlations, and applied edge-quantile thresholding (*q* = 0.65) (sensitivity analyses appear in Supplementary Table S4) to obtain binary adjacency matrices. Because there were two possible change-points, we ran PARROT in multiple-CP mode. Across real-data analyses, visualizations were generated with `ggplot2, ggraph/tidygraph`, and `ggalluvial,` and GO enrichment annotations used `clusterProfiler` (Wickham, 2016; Brunson, 2020; Yu et al., 2012).

## 3. Results

### 3.1 PARROT overview

PARROT takes as input a time-ordered sequence of networks represented as binary or weighted adjacency matrices from either unipartite or bipartite networks along with a specified number of communities *Q* (Figure 1). For each candidate change-point, PARROT partitions the series into pre- and post-transition segments, fits separate Stochastic Block Models (SBMs) via Variational Expectation–Maximization (VEM), and computes segment-wise Evidence Lower Bounds (ELBOs). The optimal change-point is identified as the split maximizing the summed profile log-likelihood across segments. Statistical significance is assessed through a permutation-based likelihood ratio test, which compares the observed improvement in fit against a null distribution obtained by shuffling time indices. Outputs include the detected change-point location with confidence intervals, community membership assignments for each phase, and estimated block connectivity parameters 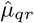 that characterize within- and between-community edge probabilities. This framework allows identification of not only *when* network structure changes but also *which* modules undergo rewiring.

**Figure 1.**
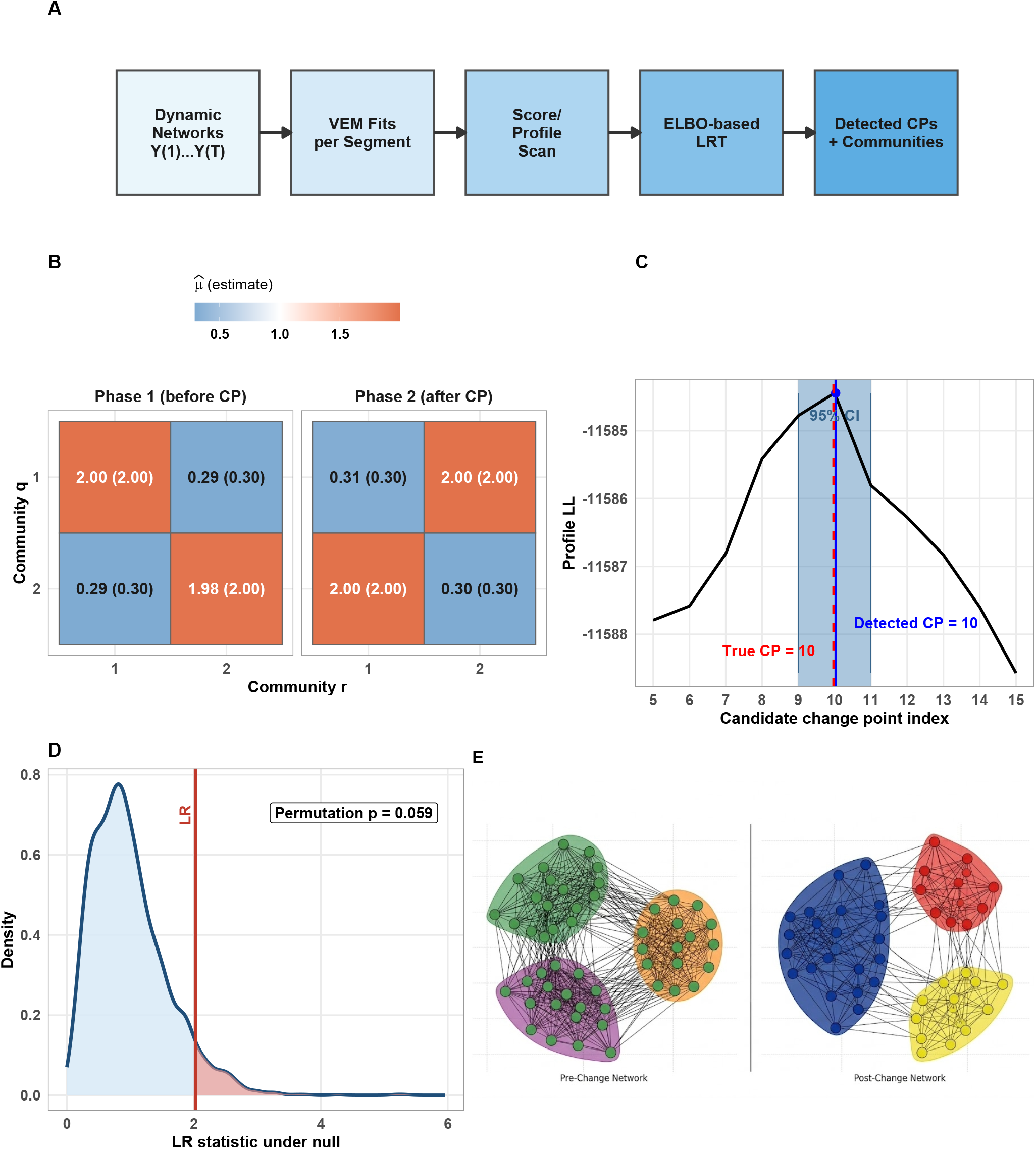
PARROT method overview. (A) Pipeline schematic: input dynamic networks are processed through VEM-based SBM fitting, score or profile scanning, ELBO-based likelihood ratio testing, yielding detected change-points with community assignments. (B) Illustration of VEM estimated SBM block means 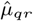 for the two phases, with true simulation values shown in parentheses. (C) Plot of the profile log-likelihood curve with detected change-point (blue solid), true change-point (red dashed), and a narrow symmetric 95% confidence interval centered on the detected split; candidate splits are scanned over indices 5–15 due to the minimum-segment constraint. (D) The permutation-null density of the likelihood-ratio (LR) test statistic, with observed LR (red line); the shaded right-tail area corresponds to the empirical permutation p-value. (E) Illustrative pre/post network visualization highlighting module-level rewiring.

### 3.2 Simulation benchmark

We first confirmed PARROT’s accuracy in recovering SBM parameters by simulating four supported network classes under static (no change-point) conditions with larger networks (Supplementary Figure S1). We then evaluated change-point detection performance against state-of-the-art methods across these network types (Table 1). To avoid ceiling effects and stress model fitting, we used smaller and shorter settings: *N* = 12 nodes (unipartite) or 8 × 10 nodes (bipartite), *T* = 10 time points, *Q* = 2 communities, true change-point at *t* = 5, and *n* = 20 replicates per setting. We simulated three scenario families: (i) community-swap transitions with partial membership rewiring, (ii) weak global mean shift with fixed community assignments, and (iii) mixed transitions combining moderate global shift with partial rewiring. Change-point simulation parameters are listed in Supplementary Table S1 and evaluation metrics are listed in Supplementary Table S2.

**Table 1.**
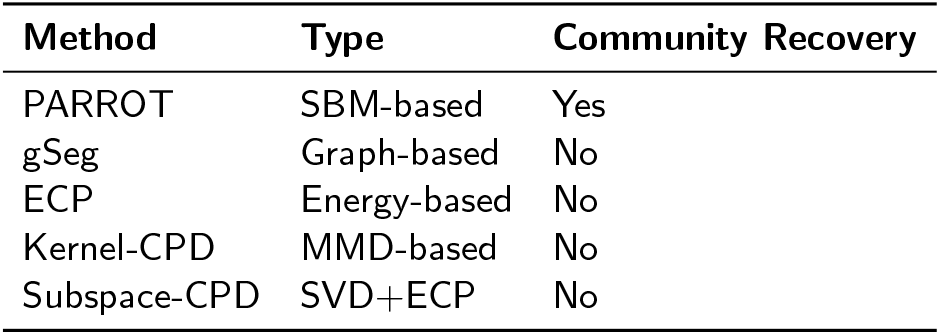
Methods compared in the simulation study.

| Method | Type | Community Recovery |
| --- | --- | --- |
| PARROT | SBM-based | Yes |
| gSeg | Graph-based | No |
| ECP | Energy-based | No |
| Kernel-CPD | MMD-based | No |
| Subspace-CPD | SVD+ECP | No |

PARROT demonstrated strong performance across all simulated scenarios and network types. Detection rates (percentage of replicates with at least one change-point detected within ±2 time points of the true boundary) showed clear separation across transition scenarios (Figure 2A): in community-swap and mixed scenarios, PARROT achieved 85– 95% detection compared with 20–30% for gSeg and 50– 70% for ECP; in global mean shift scenarios, detection rates decreased for all methods and performance gaps narrowed. F1 scores, which balance precision and recall, confirmed PARROT’s advantage across the four network types; for example, for unipartite Gaussian networks, PARROT achieved F1≈0.85 compared with *<*0.50 for gSeg (Figure 2B). Localization precision, measured by mean absolute error (MAE), showed that PARROT consistently achieved MAE*<*1 time point across community-swap and mixed scenarios, whereas competing methods often exceeded MAE*>*2 (Figure 2C). Exact-match rates revealed that PARROT most frequently recovered the true boundary at *t* = 5, achieving up to 80% exact detection in global mean shift versus 0–20% for competing methods; notably, ECP’s exact detection dropped substantially due to systematic one-time-point offsets (Figure 2D). Detailed per-method metrics are reported in Supplementary Tables S7–S19.

**Figure 2.**
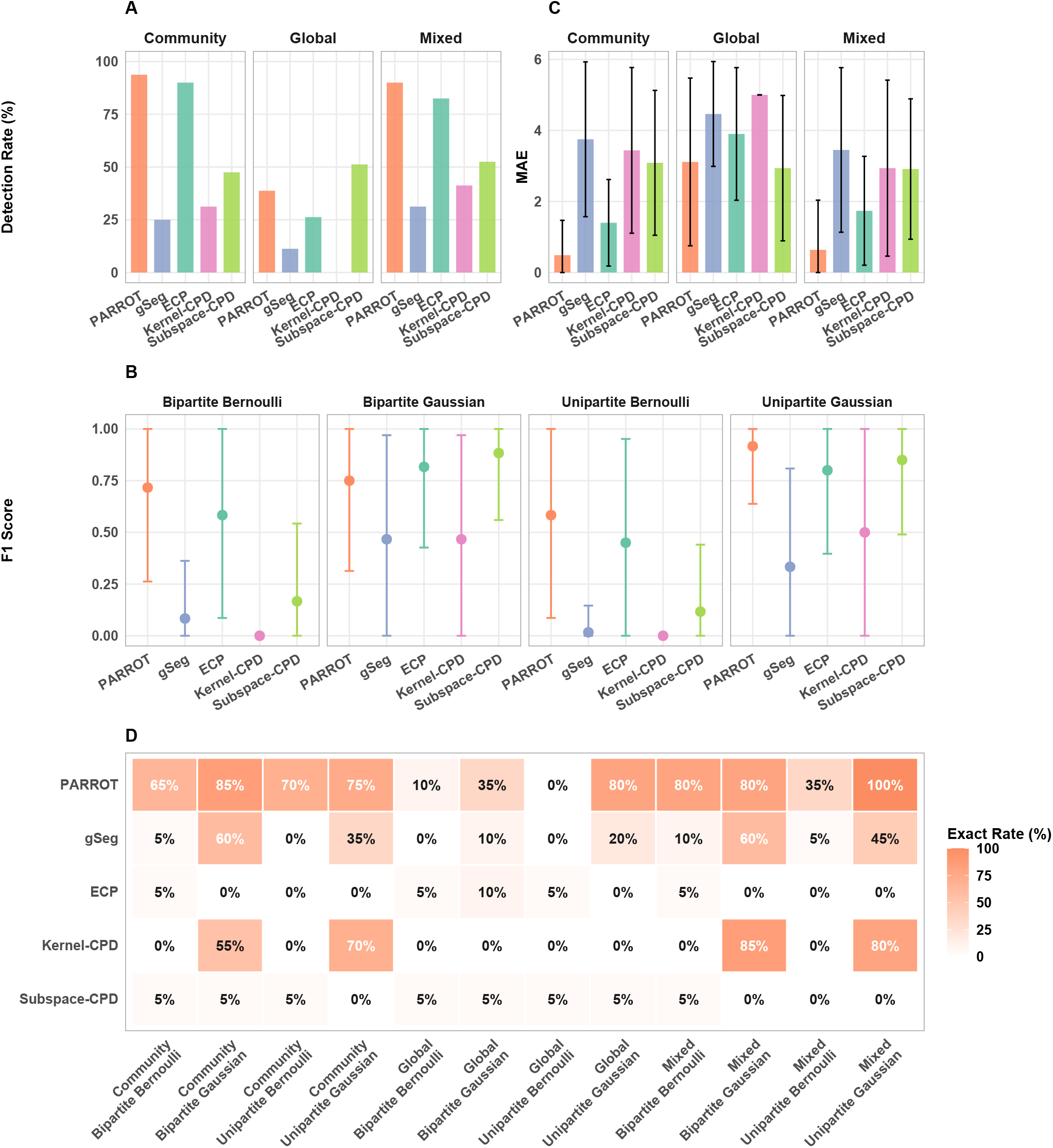
Simulation study. (A) Detection rate (percentage of replicates with at least one detected change-point within *±*2 time points) across global, community-swap, and mixed transition scenarios. (B) F1 score (mean *±* s.d. over replicates) stratified by network type, computed from precision and recall of detected versus true change-points. (C) Mean absolute error (MAE; average distance between detected and true change-point indices) by scenario; lower values indicate more precise localization. (D) Exact detection rate heatmap (percentage of replicates recovering the true change-point at *t* = 5) across method–scenario–network combinations. Simulation settings: unipartite *N* = 12, bipartite 8 *×* 10, *T* = 10, true CP *t* = 5, *n* = 20 replicates per configuration; detailed scenario tables in Supplementary Tables S8–S19.

### 3.3 Human cardiac differentiation TF–gene networks

To evaluate PARROT on real biological data, we analyzed GSE202398 (Galdos et al., 2023; Omnibus, 2023), a hiPSC cardiac differentiation scRNA-seq dataset spanning 12 time points (Day0–Day7, Day11, Day13, Day15, Day30) across two cell lines (WTC and SCVI111) (Figure 3A). Following demultiplexing cells by hashtag oligos and performing per-sample quality control, we constructed pooled bipartite TF→gene networks per day by averaging Fisher-*z*-transformed cell line-specific Spearman TF–target associations (*N*_1_ = 25 TFs, *N*_2_ = 150 target genes) (See Methods for additional details).

**Figure 3.**
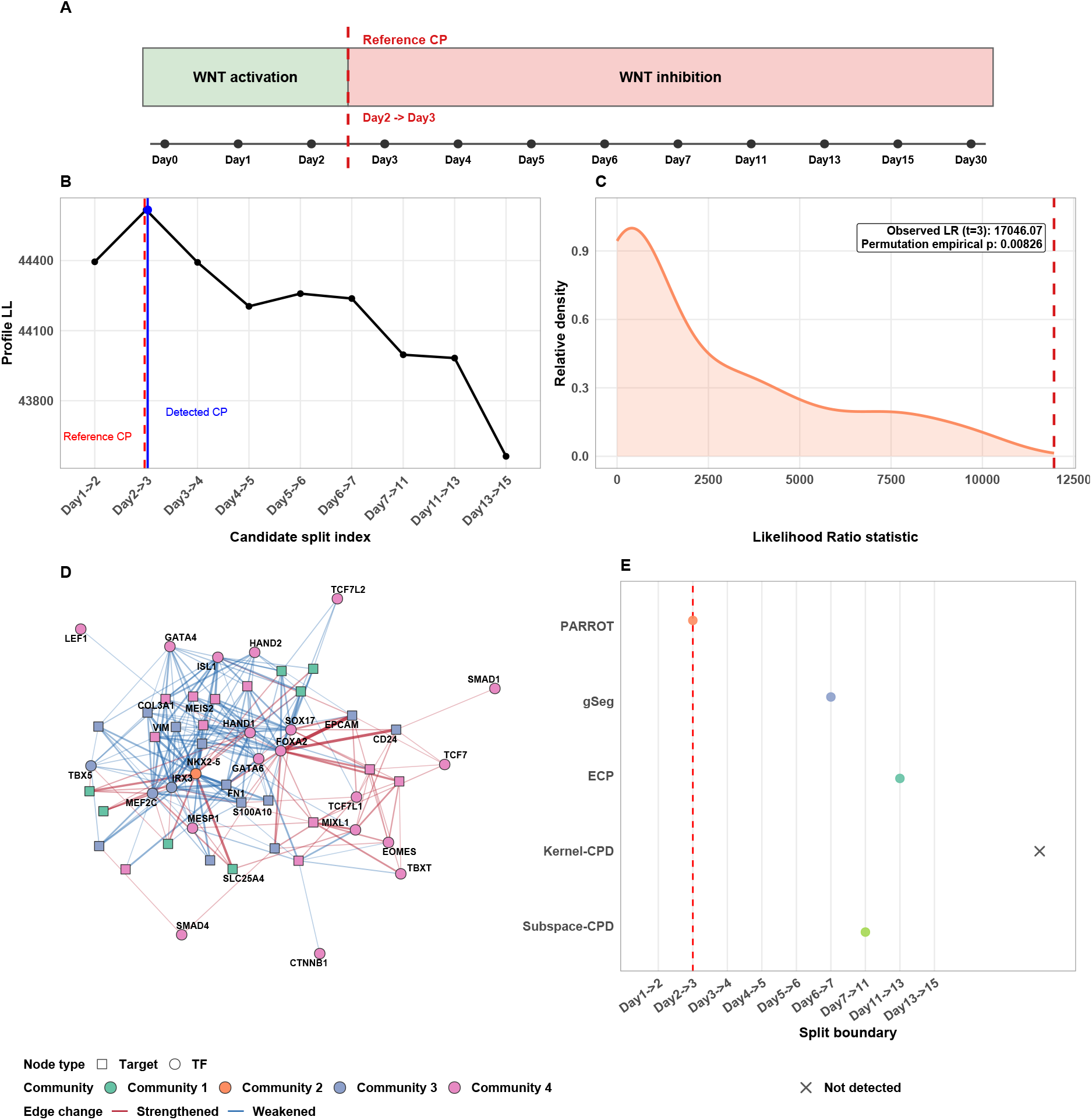
Application of change-point detection to hiPSC cardiac differentiation TF–gene networks. (A) Experimental design across 12 days with protocol phases (WNT activation then inhibition) and protocol-defined reference boundary Day2*→*Day3 (red dashed). (B) Profile log-likelihood in single-CP mode with detected CP (blue solid) and protocol reference CP (red dashed). (C) LR inference at the detected CP using a global full-series permutation null, with observed LR (dashed vertical line) and shaded right-tail empirical p-value area. (D) Gene Regulatory Network before and after the detected CP, with edges colored to highlight rewiring between phases (red = strengthened, blue = weakened) and nodes filled by their post-CP community assignment (color legend); built from top rewired targets after dropping isolates and retaining the largest connected component. (E) CP detection across five methods (PARROT, gSeg, ECP, Kernel-CPD, Subspace-CPD) against the protocol-defined reference boundary, with all methods constrained to single-CP output for fairness.

We used PARROT in single-CP mode with *Q* = 4 communities (selected by ICL; Supplementary Figure S2). PARROT identified a change-point at Day2→Day3, matching the protocol-defined WNT switch boundary (Figure 3B) (Galdos et al., 2023). This change-point was statistically significant under a full-series permutation null (Permutation *p* = 0.008; Figure 3C). Figure 3D shows the strongest rewiring events (top rewired targets, edge changes above the 70th percentile, largest component only), revealing biologically coherent shifts: WNT-axis regulators (TCF/LEF, SMAD) alter connectivity with mesoderm markers (MESP1, EOMES) and early cardiac genes (GATA4, NKX2-5), consistent with the Day2→Day3 transition from WNT activation to inhibition.

Comparison across five methods showed that PARROT was the only method to identify the protocol-defined Day2→Day3 boundary when all methods were constrained to single-CP output (Figure 3E); competing methods detected change-points at various times and none recovered the true transition.

### 3.4 Mouse postnatal lung development co-expression networks

To assess PARROT’s performance on unipartite co-expression networks with multiple candidate boundaries, we used GEO series GSE74243 (Beauchemin et al., 2016; Omnibus, 2023), which profiles postnatal mouse lung transcriptomes across three strains during early development; Figure 4A shows seven pooled time points aligned to the original molecular staging. We constructed unipartite co-expression networks by computing pairwise Spearman correlations among the top 100 variable genes, thresholding at quantile *q* = 0.65 to obtain binary adjacency matrices. Model selection favored *Q* = 6 communities based on ICL diagnostics (Supplementary Figure S2). We then ran PARROT in multiple-CP mode (max *K* = 2) to test whether we could identify the biologically defined reference boundary from the original study, T5→T6 (ALV4→MAT) and to test whether additional within-alveolarization candidate transitions are detectable.

**Figure 4.**
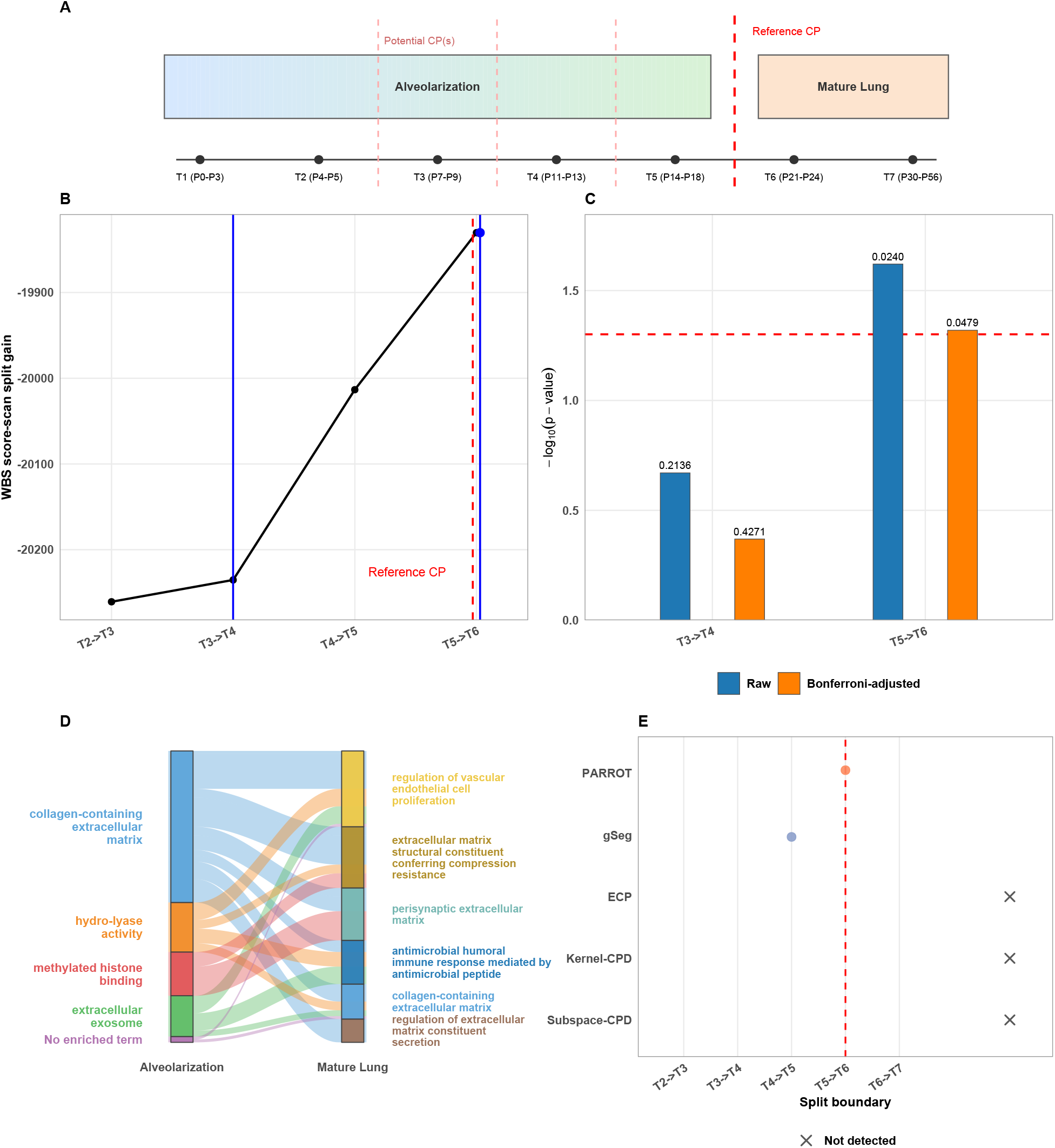
Application to mouse postnatal lung development co-expression networks. (A) Seven-time-point design aligned to postnatal staging, with one reference boundary of red dashed line (*T* 5 *→ T* 6) and lighter dashed lines marking additional candidate boundaries (*T* 2 *→ T* 3, *T* 3 *→ T* 4, *T* 4 *→ T* 5). (B) WBS score-scan split gain across feasible boundaries, labeled as *T*_*i*_ *→ T*_*i*+1_. With min segment = 2 and *T* 5 *→ T* 6 selected first by gain, *T* 4 *→ T* 5 is excluded by spacing and *T* 3 *→ T* 4 is selected as the second candidate. (C) Direct per-CP permutation significance for detected CPs using permutation tests (*n*_perm_ = 500), shown as grouped raw and Bonferroni-adjusted *−* log_10_(*p*) bars; *T* 5 *→ T* 6 has the stronger raw signal and remains significant after Bonferroni adjustment. (D) Two-stage Sankey across the *T* 5 *→ T* 6 transition (Alveolarization*→*Mature Lung), with strata labeled by functional enrichment terms selected from GO (BP/CC/MF) using all genes per module. (E) CP detection across five methods (PARROT, gSeg, ECP, Kernel-CPD, Subspace-CPD) against the single reference boundary *T* 5 *→ T* 6.

PARROT identified T5→T6 as its top candidate change-point, consistent with biological expectations. In the multiple-CP WBS filtering step, we use min segment = 2. Because T5→T6 was selected first (largest split gain), the adjacent split at T4→T5 is excluded by the spacing constraint, and T3→T4 was selected as a second candidate. Under direct per-CP global permutation testing (*n*_perm_ = 500), T5→T6 showed the stronger raw signal (raw *p* = 0.024) than T3→T4 (raw *p* = 0.214), and T5→T6 remained significant after Bonferroni adjustment (adjusted *p* = 0.048; Figure 4B–C).

Given the strong statistical support for T5→T6, we characterized module rewiring across this transition using a two-stage Sankey diagram (Alveolarization→Mature Lung; Figure 4D). Functional enrichment of module genes using `clusterProfiler` (GO BP/CC/MF) revealed substantial overlap between ALV and MAT modules, consistent with continuity between late alveolarization and early mature homeostasis. This pattern indicates rewiring within shared functional programs—particularly extra-cellular matrix (ECM) remodeling and vascular development associated with postnatal septation and capillary maturation (Beauchemin et al., 2016)— rather than wholesale replacement of module function. In method comparison, PARROT detected the T5→T6 while gSeg found the secondary T4→T5 and the other methods failed to identify a change-point (Figure 4E).

## 4. Discussion

### 4.1 Main findings

PARROT addresses a critical gap in network change-point detection: the joint estimation of transition timing and community structure. Although several competing methods perform reasonably well in terms of change-point *detection* accuracy in some settings (Figure 2), the comparison is not apples-to-apples because the methods are detecting fundamentally different signals. A two-stage workflow—apply a change-point method to a network summary and then run Louvain clustering on each resulting segment—selects change points that optimize a global topological summary (such as edge count, kernel distance, spectral statistic) and only describes communities *after the fact*. PARROT instead selects the change-point that *best reflects a change in community structure*: the profile objective rewards splits at which segment-specific SBMs jointly fit the observed snapshots better than a single shared SBM. This shift is more than wording; it lets PARROT recover the change-point that captures the cell’s transition between distinct modular regulatory programs (Kashtan and Alon, 2005), for example, when the cell reorganizes which genes serve which regulatory module. This interpretability—identifying the modules whose connectivity is rewired —is essential for translational applications where researchers need actionable targets. In this sense, PARROT directly speaks to our intuition about changes in biological state being linked to alterations in regulatory network community structure to activate or repress specific biological functions.

The two biological applications highlight PARROT’s ability to operate across distinct network architectures while delivering interpretable results. In the cardiac differentiation study (bipartite Gaussian; Figure 3), PARROT not only recovered the protocol-defined Wnt-switching boundary at Day2→Day3 but also revealed the specific TF–gene modules undergoing rewiring—information inaccessible to methods that treat networks as unstructured objects. The mouse lung development analysis (unipartite binary; Figure 4) further demonstrated PARROT’s capacity for multiple change-point inference, pinpointing T5→T6 as the statistically dominant transition even after Bonferroni correction and linking it to ECM remodeling and vascular maturation programs.

### 4.2 Limitations and future directions

PARROT requires specification of the number of communities *Q* although it supports ICL-based model selection to identify a likely number of communities. The method also assumes an underlying block structure with reasonably distinct modules, which may not hold for all networks. By design, PARROT detects structural changes in community connectivity rather than global mean shifts; consequently, scenarios involving only weak homogeneous signal changes fall outside its intended scope. For exploratory analyses where community structure is not required, distribution-free methods such as gSeg remain excellent alternatives.

As in all change-point methods, estimation variance increases near the beginning and end of the time-course sequence, where one segment is necessarily short (Andrews, 1993; Bai and Perron, 1998). This issue is amplified when *T* is small and WBS recursively subdivides the series (Fryzlewicz, 2014), because fitting an SBM with *Q* communities requires estimating *O*(*Q*^2^) block parameters (Daudin et al., 2008; Celisse et al., 2012); if the resulting segments are too short relative to *Q*, parameter estimates and community assignments become unstable. PARROT’s permutation test partially mitigates this problem because each permutation replicate applies the same LR construction—null SBM versus two segment-specific SBMs at a fixed split—to time-shuffled data, finite-sample fitting variability enters both the observed statistic and the reference distribution, providing better-calibrated significance assessment. Nonetheless, community assignments from very short segments should be interpreted with caution, and practitioners may need to reduce *Q* or increase temporal resolution when the number of timepoints, *T*, is limited.

PARROT assumes the same number of communities Q before and after the change point, while allowing memberships and interaction parameters to vary across segments. This preserves the nested SBM structure required for the likelihood-ratio test in Eq. (5). PARROT therefore detects two kinds of change-point: community reorganization, where genes or TFs are reassigned across modules so that module composition and size change (Figure 4D), and edge rewiring, where the same nodes connect differently across modules (in Figure 3D, similar target genes are regulated by a different set of TFs after the transition). Extending the framework to allow *Q*_1_ ≠ *Q*_2_ is left for future work.

#### Compatible regulatory-network methods

PARROT operates on a sequence of network snapshots and is agnostic to how each snapshot is constructed, provided that edge weights are approximately Gaussian or that edges are coded as Bernoulli presence/absence after thresholding. Methods that fit naturally into the Gaussian variant after standard transformations include Pearson/Spearman co-expression with Fisher-*z* on the correlations, WGCNA (Langfelder and Horvath, 2008) after soft-thresholding (treat the topological-overlap matrix entries as weights), ARACNe mutual-information networks (Margolin et al., 2006) after rank or log transformation of the MI scores, PANDA TF→gene message-passing scores (Glass et al., 2013), Inferelator regression weights (Bonneau et al., 2006) after Fisher-*z*, and CLR/GENIE3 importance scores (Huynh-Thu et al., 2010). SCENIC’s regulon AUCell matrices (Aibar et al., 2017) are best handled in the Bernoulli variant after binarizing AUCell activity.

#### Scalability and the small-N regime

The biological time series networks we analyzed deliberately use small node counts (*N* = 25 × 150 for the cardiac bipartite TF–gene network and *N* = 100 for the lung co-expression network) and short sequences (*T* ≤ 12), reflecting the typical resolution of public time-resolved GEO datasets. PARROT’s per-iteration VEM cost is *O*(*T NQ*^2^) per segment and WBS adds *O*(*M*) random-interval scans; memory is *O*(*N* ^2^*T*) for the cumulative sufficient statistics. Empirically, end-to-end runs on *N* ∼ 200, *T* ∼ 20 complete in seconds-to-minutes on a laptop; for *N* ∼ 10^3^ we recommend the score-scan mode (Supplementary Table S6) and consider sparse-edge thresholding before fitting. Very large regulatory networks (*N* ≫ 10^3^) will benefit from degree-corrected SBMs and stochastic-VEM updates, which we leave as future work.

## Acknowledgements

The authors thank Viola Fanfani, Katherine H. Shutta, Camila Lopes-Ramos, Marouen Ben Guebila, and Jonas Fischer for discussions regarding the method.

## Author contributions

CC developed the PARROT package, implemented analyses, and drafted the manuscript. CC, MP, and JQ conceived the study and revised the manuscript.

## Supplementary data

Supplementary data are available at *Bioinformatics* online.

## Conflict of interest

The authors declare no competing interests.

## Funding

CC and JQ were supported by funding from the US National Cancer Institute (R35CA220523) and the National Human Genome Research Institute (R01HG011393). MP was supported by the US National Cancer Institute (R01CA251729).

## Data availability

PARROT is available at https://github.com/cchen22/ PARROT and https://github.com/netZoo/netZooR (Ben Guebila et al., 2023). Processed outputs used to generate manuscript figures are included in this repository; source GEO accession identifiers for real-data analyses are reported in the Methods section.

## Notes

https://github.com/netZoo/netZooR

### Competing Interest Statement

The authors have declared no competing interest.

