## Supplementary Material for "PARROT: Phase-Altering Regulatory Rewiring Over Time"

### 1 Supplementary Methods

#### 1.1 Variational EM Algorithm

We fit a stochastic block model (SBM) (Holland et al., 1983) to each network segment by Variational EM. Let $\mathbf{Y} = \{Y^{(t)}\}_{t=1}^{T}$ denote the observed adjacency (or weight) tensor, $\mathbf{Z}$ the latent block memberships, and $\theta = (\boldsymbol{\pi}, \boldsymbol{\Phi})$ the model parameters, where $\boldsymbol{\pi}$ collects the mixing proportions and $\boldsymbol{\Phi}$ the block-level parameters ($\mu_{qr}$ and $\sigma_{qr}$ for Gaussian, $p_{qr}$ for Bernoulli).

The variational distribution factorizes over node memberships. For unipartite networks a single set of variational parameters $\tau_{iq}$ ($i = 1, \ldots, N$; $q = 1, \ldots, Q$) is maintained with mixing proportions $\pi_q$ ($q = 1, \ldots, Q$). For bipartite networks two independent sets $\tau_{iq}^{(1)}$ ($i = 1, \ldots, N_1$; $q = 1, \ldots, Q_1$) and $\tau_{jl}^{(2)}$ ($j = 1, \ldots, N_2$; $l = 1, \ldots, Q_2$) are maintained with separate mixing proportions $\pi_q^{(1)}$ and $\pi_l^{(2)}$. Mixing proportions are initialized uniformly ($\pi_q = 1/Q$); $\tau$ is initialized from random hard assignments, and block parameters from random draws (see below). Each iteration cycles: E-step $\to$ M-step $\to$ ELBO evaluation.

**E-step.** Variational parameters are updated in log-space followed by softmax normalization. For Gaussian networks:

$$\log \tau_{iq}^{\text{new}} = \log \pi_q + \sum_r \sum_j \tau_{jr} \left[ -\tfrac{T}{2} \log(2\pi\sigma_{qr}^2) - \tfrac{1}{2\sigma_{qr}^2} \left( S_{ij} - 2\mu_{qr} Y_{ij}^{\Sigma} + \mu_{qr}^2 T \right) \right] \tag{1}$$

For Bernoulli networks:

$$\log \tau_{iq}^{\text{new}} = \log \pi_q + \sum_r \sum_j \tau_{jr} \left[ n_{ij} \log p_{qr} + (T - n_{ij}) \log(1 - p_{qr}) \right] \tag{2}$$

where $S_{ij} = \sum_t Y_{ij}^{(t)2}$, $Y_{ij}^{\Sigma} = \sum_t Y_{ij}^{(t)}$, and $n_{ij} = \sum_t Y_{ij}^{(t)}$. Here $\pi_q$, $\mu_{qr}$, $\sigma_{qr}$, and $p_{qr}$ denote current-iteration values from the preceding M-step (or initialization at $k$=0). In the unipartite case the inner sum runs over $j \neq i$; in the bipartite case it runs over all $N_2$ column nodes using the current $\tau^{(2)}$, then $\tau^{(2)}$ is updated by summing over all $N_1$ row nodes using the refreshed $\tau^{(1)}$.

**M-step.** Mixing proportions (with subsequent renormalization):

$$\hat{\pi}_q = \frac{1}{N} \sum_{i=1}^{N} \tau_{iq} \tag{3}$$

(one set for unipartite; for bipartite, $\hat{\pi}_q^{(1)} = N_1^{-1} \sum_i \tau_{iq}^{(1)}$ and $\hat{\pi}_l^{(2)} = N_2^{-1} \sum_j \tau_{jl}^{(2)}$ separately). Block parameters use expected pair weights $w_{ij}^{qr} = \tau_{iq}\tau_{jr}$ (unipartite) or $w_{ij}^{ql} = \tau_{iq}^{(1)}\tau_{jl}^{(2)}$ (bipartite):

$$\text{Gaussian mean:} \quad \hat{\mu}_{qr} = \frac{\sum_{i,j} w_{ij}^{qr} Y_{ij}^{\Sigma}}{\sum_{i,j} w_{ij}^{qr} \cdot T} \tag{4}$$

$$\text{Bernoulli probability:} \quad \hat{p}_{qr} = \frac{\sum_{i,j} w_{ij}^{qr} n_{ij}}{\sum_{i,j} w_{ij}^{qr} \cdot T} \tag{5}$$

For Gaussian networks a noise scale is additionally estimated. Under the default shared-variance model (`block_sigma = FALSE`):

$$\hat{\sigma} = \max\left(0.01, \sqrt{\frac{\sum_{q,r} \sum_{i,j} w_{ij}^{qr} \sum_t (Y_{ij}^{(t)} - \hat{\mu}_{qr})^2}{\sum_{q,r} \sum_{i,j} w_{ij}^{qr} \cdot T}}\right) \tag{6}$$

With per-block variances (`block_sigma=TRUE`), each $\hat{\sigma}_{qr}$ is estimated analogously from the $(q, r)$-specific weighted residuals. For unipartite networks, all block-parameter matrices are symmetrized and edge sums run over the upper triangle ($i < j$).

**ELBO.** The Evidence Lower Bound decomposes as:

$$\mathcal{L} = \underbrace{\textstyle\sum_{i,q} \tau_{iq} \log \pi_q}_{\text{mixing prior}} + \underbrace{\mathbb{E}_{\mathcal{Q}}[\log p(\mathbf{Y} \mid \mathbf{Z}; \mathbf{\Phi})]}_{\text{expected log-likelihood}} - \underbrace{\textstyle\sum_{i,q} \tau_{iq} \log \tau_{iq}}_{\text{entropy}} \tag{7}$$

For bipartite networks the mixing-prior and entropy terms each include both the row and column sides. For unipartite networks the expected log-likelihood is computed over the upper triangle ($i < j$) and doubled by symmetry.

**Convergence and initialization.** The algorithm is run from multiple random initializations; the run with the highest final $\mathcal{L}$ is retained. Convergence is declared when the absolute ELBO change $|\mathcal{L}^{(k)} - \mathcal{L}^{(k-1)}|$ falls below tolerance $\epsilon$ (default $10^{-6}$).

#### 1.2 ICL Model Selection

The Integrated Classification Likelihood (ICL) penalizes the ELBO by the number of free parameters under standard BIC-type scaling:

$$\text{ICL}(Q) = \text{ELBO} - \underbrace{\tfrac{d_{\text{mix}}}{2} \log n_{\text{mix}}}_{\text{mixing-proportion penalty}} - \underbrace{\tfrac{d_{\text{block}} + d_\sigma}{2} \log n_{\text{obs}}}_{\text{block-parameter penalty}} \tag{8}$$

The parameter counts and effective sample sizes depend on the network type:

| | **Unipartite** | **Bipartite** |
|---|---|---|
| Block params $d_{\text{block}}$ | $Q(Q+1)/2$ | $Q_1 Q_2$ |
| Mixing params $d_{\text{mix}}$ | $Q - 1$ | $(Q_1 - 1) + (Q_2 - 1)$ |
| Mixing scale $n_{\text{mix}}$ | $N$ | $\max(N_1, N_2)$ |
| Observation scale $n_{\text{obs}}$ | $N(N-1)T/2$ | $N_1 N_2 T$ |

For Gaussian networks an additional variance penalty $d_\sigma$ is included: $d_\sigma = 1$ when a single shared variance is estimated, or $d_\sigma = d_{\text{block}}$ when per-block variances are used (`block_sigma=TRUE`). For Bernoulli networks $d_\sigma = 0$. The community number $Q$ (or the pair $Q_1, Q_2$) that maximizes ICL$(Q)$ is selected.
For both network types, $n_{\text{obs}}$ equals the number of distinct dyads times $T$: $N(N-1)T/2$ for undirected unipartite networks (Daudin et al., 2008) and $N_1 N_2 T$ for bipartite networks.

#### 1.3 Wild Binary Segmentation

Algorithm 1 describes the WBS procedure for multiple change point detection.

**Algorithm 1:** Wild Binary Segmentation for Multiple Change Points

```
Input: Network sequence Y, minimum segment m, number of random intervals M, maximum
       CPs K
Output: Filtered change-point set C
Initialize candidate list A ← ∅;
for j = 1, ..., M do
    Sample random interval [s_j, e_j] with length ≥ 2m + 1;
    Fit interval null model and compute ELBO_0 on [s_j, e_j];
    for each valid split τ ∈ [s_j + m, e_j − m] do
        Fit left/right segment models and compute gain;
        g(τ) = ELBO_L(τ) + ELBO_R(τ) − ELBO_0;
    end
    τ̂_j ← arg max_τ g(τ);
    if g(τ̂_j) > 0 then
        Add candidate (τ̂_j, g(τ̂_j)) to A;
    end
end
Aggregate duplicate candidate positions by keeping the maximum gain;
Sort unique candidates by decreasing gain;
Initialize C ← ∅;
for candidate τ in sorted order do
    if τ is at least m away from all CPs already in C then
        Add τ to C;
    end
    if |C| = K then
        break;
    end
end
return sorted C
```

**Score scan variant (default).** Algorithm 1 describes the *profile* variant, which refits the SBM at every candidate split. PARROT defaults to a faster *score scan*: a single global SBM is fitted on the full sequence and the resulting variational memberships are held fixed, so that cumulative sufficient statistics yield closed-form block log-likelihoods for any candidate segment. The gain at each split is the two-segment block log-likelihood minus the single-segment value, replacing the ELBO difference used by the profile variant. Supplementary Figure S3 compares the runtime–accuracy tradeoff between the two scan modes.

**Current inferential limitation.** The multiple-CP procedure above is a practical candidate-generation and filtering strategy, but it does not currently provide formal global Type-I error control across all detected boundaries. Single-CP p-values are therefore reported using permutation/bootstrap nulls and should be interpreted as approximate under possible temporal dependence.

### 2 Supplementary Results

#### 2.1 SBM Parameter Recovery

Before evaluating change point detection, we verified that PARROT's Variational EM correctly recovers SBM parameters under static (no change point) conditions. Figure S1 shows results across all four network types—unipartite Gaussian, unipartite Bernoulli, bipartite Gaussian, and bipartite Bernoulli.

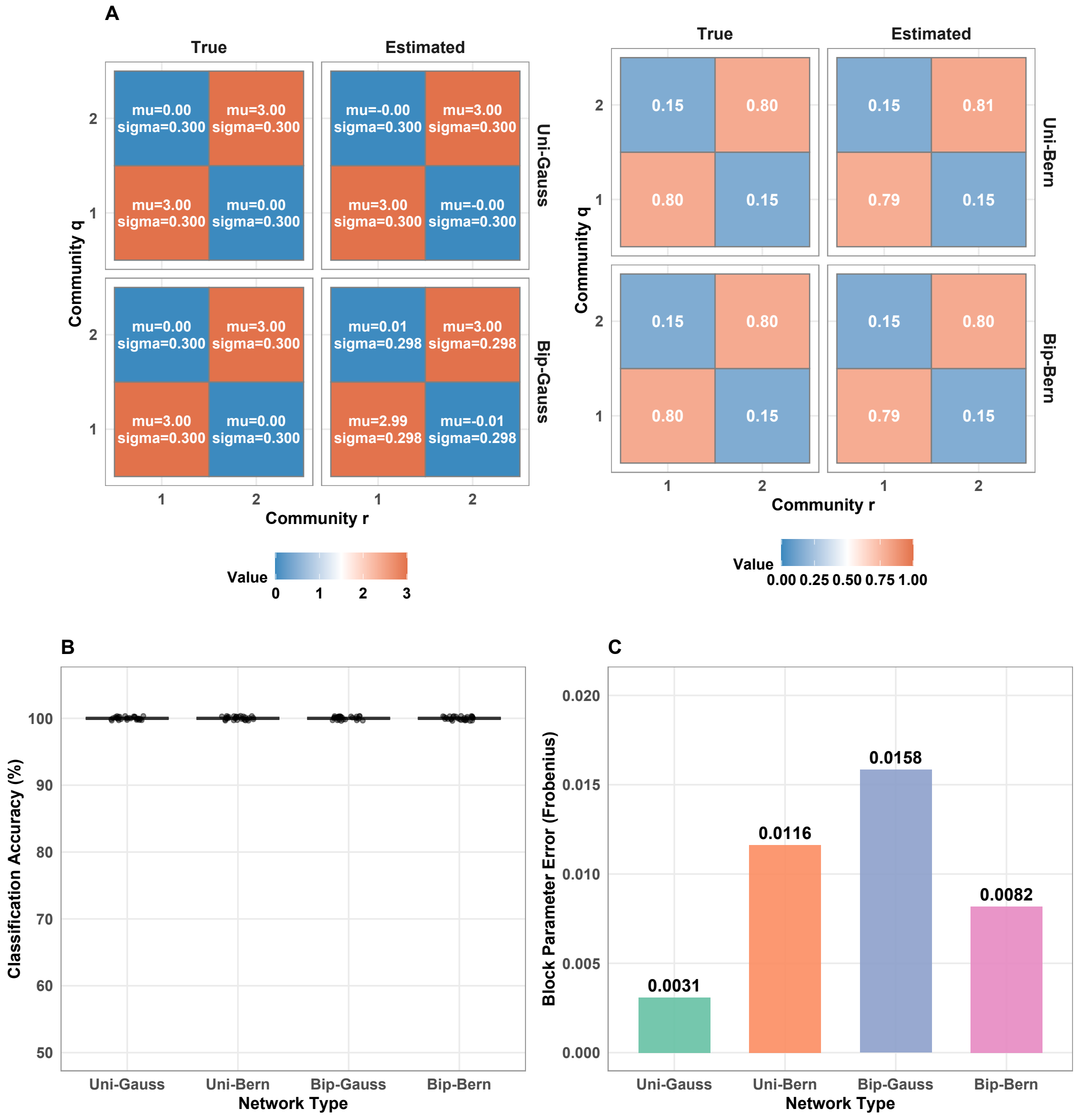

Figure S1: **SBM parameter recovery across four network types (static networks, no change point).** (A) True vs estimated block parameter matrices for unipartite and bipartite networks. Gaussian cells report both block mean ($\mu$) and noise scale ($\sigma$), while Bernoulli cells report block probability. (B) Classification accuracy (percentage of nodes correctly assigned to communities) across 30 replicates per network type. (C) Block parameter estimation error (Frobenius norm of estimated vs true block matrices) for each network type from a representative replicate. Settings: unipartite $N = 40$, bipartite $15 \times 20$, $T = 20$, $Q = 2$; these are separate from the change point benchmark settings in Table S1.

#### 2.2 Simulation Settings and Evaluation Metrics

This section documents the simulation parameters and evaluation metrics used for the change point detection benchmark (Figure 2). Table S1 summarizes data-generation settings (network sizes, true CP location, scenario definitions, and replicate counts), and Table S2 defines the evaluation metrics and matching rules used for method comparisons.

##### 2.2.1 Simulation Settings

Table S1: Simulation settings for the global, community-swap, and mixed transition comparison study (Figure 2).

| **Network Type** | $N_1$ | $N_2$ | $T$ | $\tau^*$ | $Q$ | **Replicates** |
|---|---|---|---|---|---|---|
| Unipartite Gaussian | 12 | – | 10 | 5 | 2 | 20 |
| Unipartite Bernoulli | 12 | – | 10 | 5 | 2 | 20 |
| Bipartite Gaussian | 8 | 10 | 10 | 5 | 2 | 20 |
| Bipartite Bernoulli | 8 | 10 | 10 | 5 | 2 | 20 |

| **Parameter** | **Value** |
|---|---|
| Tolerance for CP matching | $\pm 2$ time points |
| *Global Mean Shift Scenario (weak-to-moderate signal):* | |
| Gaussian noise $\sigma$ | 0.55 |
| Gaussian $\mu_{\text{base}}$ | $\begin{pmatrix} 1.8 & 0.6 \\ 0.6 & 1.8 \end{pmatrix}$ |
| Gaussian global shift | 0.25 (added to all $\mu$ entries) |
| Bernoulli $p_{\text{base}}$ | $\begin{pmatrix} 0.58 & 0.32 \\ 0.32 & 0.58 \end{pmatrix}$ |
| Bernoulli probability shift | 0.07 (added to all entries) |
| *Community Swap Scenario (structural rewiring):* | |
| Gaussian noise $\sigma$ | 0.45 |
| Gaussian $\mu_{\text{base}}$ | $\begin{pmatrix} 2.2 & 0.7 \\ 0.7 & 2.2 \end{pmatrix}$ |
| Bernoulli $p_{\text{base}}$ | $\begin{pmatrix} 0.70 & 0.30 \\ 0.30 & 0.70 \end{pmatrix}$ |
| Community swap fraction | 35% of nodes switch community |
| *Mixed Transition Scenario (global + structural):* | |
| Gaussian noise $\sigma$ | 0.50 |
| Gaussian $\mu_{\text{base}}$ | $\begin{pmatrix} 2.0 & 0.55 \\ 0.55 & 2.0 \end{pmatrix}$ |
| Gaussian global shift | 0.15 (added to all $\mu$ entries) |
| Bernoulli $p_{\text{base}}$ | $\begin{pmatrix} 0.64 & 0.30 \\ 0.30 & 0.64 \end{pmatrix}$ |
| Bernoulli probability shift | 0.05 (added to all entries) |
| Community swap fraction | 30% of nodes switch community |

**Notes:** $N_1$ = number of row nodes, $N_2$ = number of column nodes (bipartite only), $T$ = number of time points, $\tau^*$ = true change point location, $Q$ = number of communities. For community swap scenarios, block parameters remain fixed while node memberships change. For mixed scenarios, both block parameters and node memberships change.

##### 2.2.2 Evaluation Metrics

Table S2: Evaluation metrics used in the simulation study.

| **Metric** | **Description** |
|---|---|
| Precision | Proportion of detected change points that match a true change point (within tolerance): $\frac{\text{TP}}{\text{TP+FP}}$ |
| Recall | Proportion of true change points that are detected (within tolerance): $\frac{\text{TP}}{\text{TP+FN}}$ |
| F1 Score | Harmonic mean of precision and recall: $\frac{2\times\text{Precision}\times\text{Recall}}{\text{Precision}+\text{Recall}}$ |
| MAE | Mean Absolute Error: average distance (in time points) between each detected CP and its matched true CP |
| Detection Rate | Same as Recall; proportion of true CPs successfully detected |
| Exact Rate | Proportion of true CPs detected at exactly the correct time point (MAE = 0) |

**Note:** A detected change point is considered a true positive (TP) if it falls within $\pm 2$ time points of a true change point. Each true CP can match at most one detected CP.

#### 2.3 Real-data community-number selection (Q)

We evaluated community numbers on a compact grid ($Q = 1, \ldots, 6$) for both real-data applications using the ICL criterion computed from full-series SBM fits (all snapshots jointly). Dashed red lines indicate ICL-optimal $Q$. For the human cardiac differentiation dataset (GSE202398 (Galdos et al., 2023)), the Q-scan uses the bipartite regulatory network (25 TFs, 150 targets; Gaussian SBM with `block_sigma=FALSE`), for which the ICL optimum coincides with $Q = 4$. For the mouse lung development dataset (GSE74243 (Beauchemin et al., 2016)), the Q-scan uses the unipartite co-expression network (100 genes; Bernoulli SBM), for which the ICL optimum coincides with $Q = 6$.

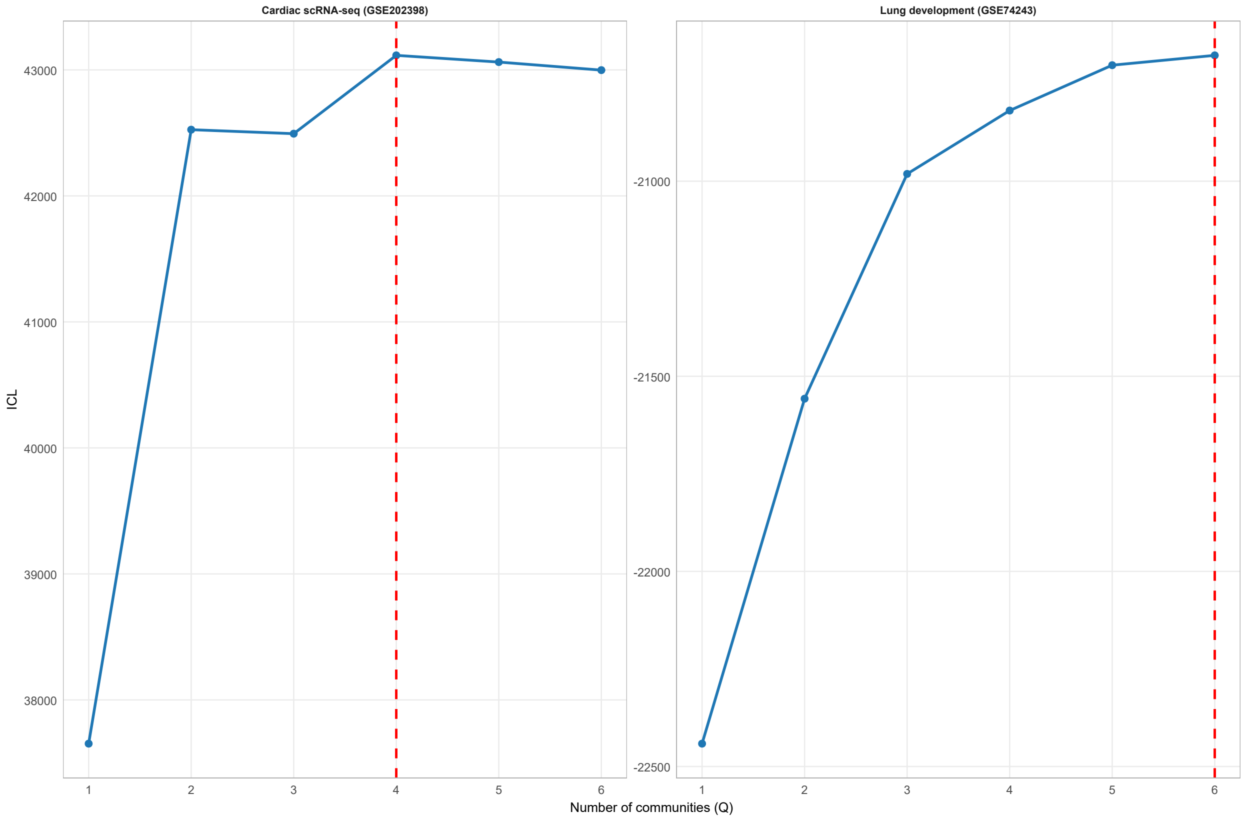

Figure S2: **Supplementary Figure S2. Real-data selection of community number $Q$.** Community-number scans for the human cardiac scRNA-seq bipartite network (GSE202398) and the mouse postnatal lung development network (GSE74243) using ICL. Dashed red lines indicate ICL-optimal $Q$.

#### 2.4 Score scan versus profile-likelihood scan

We benchmark score scan against full profile-likelihood scanning to quantify the runtime–accuracy tradeoff. For each configuration/replicate, both methods are evaluated on the same simulated realization to ensure a paired comparison. In the unipartite Gaussian setting, we additionally include a stress-test with strong post-change latent-label reassignment at an early change point (20% of sequence length), while holding block parameters fixed; this setting highlights the value of segment-wise profile refitting over the score approximation.

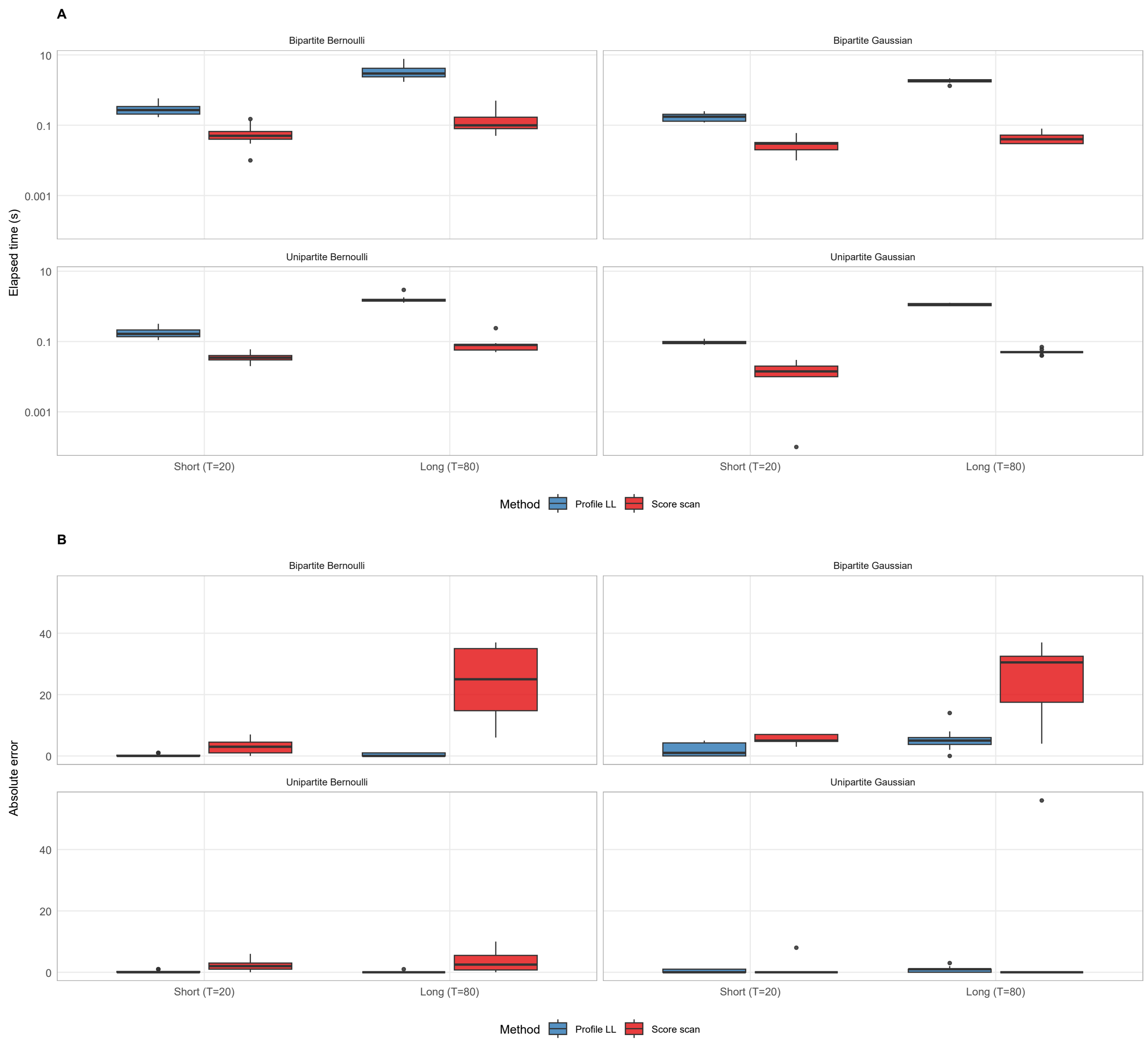

Figure S3: **Supplementary Figure S3. Runtime–accuracy comparison of score scan vs profile-likelihood scan.** Single-CP simulations across all four network classes (unipartite/bipartite $\times$ Gaussian/Bernoulli) under short ($T = 20$) and long ($T = 80$) sequences, with paired method comparisons on identical simulated networks. Panel A shows elapsed runtime (log scale); Panel B shows localization error $|\hat{\tau} - \tau_{\text{true}}|$. The unipartite Gaussian facet uses a deliberate latent-label rewiring stress setting with an early change point, where profile-likelihood scanning can deliver substantially better change-point localization at higher runtime cost.

#### 2.5 Real-data preprocessing sensitivity checks

To assess robustness of real-data conclusions to preprocessing choices, we ran a compact sensitivity panel with pre-specified perturbations. For cardiac differentiation data (GSE202398), we varied selected TF and gene counts and selected community number $Q$ by ICL. For lung development data (GSE74243), we varied selected gene count, edge quantile, and selected community number $Q$ by ICL while retaining the same Beauchemin-aligned 7-time-point design (ALV1–4 + MAT with sub-splits of ALV3 and MAT).

Table S3 summarizes cardiac sensitivity under three TF/target panel sizes with ICL-selected $Q$. Table S4 summarizes lung sensitivity under nine gene/edge settings with ICL-selected $Q$ and reports whether the reference CP 5 boundary is recovered at T5→T6.

Table S3: Cardiac differentiation sensitivity analysis across three feature-panel sizes. For each setting, the community number $Q$ is selected by ICL from $Q = 1, \ldots, 6$ and PARROT is fit on the pooled bipartite Gaussian series.

| TFs | Genes | $Q_{\text{ICL}}$ | CP | Boundary | Match Day2→3 |
|---|---|---|---|---|---|
| 15 | 100 | 5 | 3 | Day2->3 | Yes |
| 25 | 150 | 4 | 3 | Day2->3 | Yes |
| 35 | 200 | 6 | 3 | Day2->3 | Yes |

Table S4: Lung development sensitivity analysis across selected gene counts and edge quantiles. For each setting, $Q$ is selected by ICL from $Q = 1, \ldots, 6$ and PARROT is fit on binary unipartite networks.

| Genes | Edge quantile | $Q_{\text{ICL}}$ | CP | Boundary | Match T5→T6 |
|---|---|---|---|---|---|
| 100 | 0.55 | 6 | 5 | T5->T6 | Yes |
| 100 | 0.65 | 6 | 5 | T5->T6 | Yes |
| 100 | 0.75 | 6 | 5 | T5->T6 | Yes |
| 150 | 0.55 | 6 | 5 | T5->T6 | Yes |
| 150 | 0.65 | 6 | 5 | T5->T6 | Yes |
| 150 | 0.75 | 6 | 5 | T5->T6 | Yes |
| 200 | 0.55 | 6 | 5 | T5->T6 | Yes |
| 200 | 0.65 | 6 | 5 | T5->T6 | Yes |
| 200 | 0.75 | 6 | 5 | T5->T6 | Yes |

#### 2.6 GO Enrichment Summaries

Gene Ontology enrichment was performed on lung-module gene sets using `clusterProfiler`. Table S5 reports all significant GO terms (BH-adjusted $p < 0.05$) by phase and module.

Table S5: Top lung-development GO enrichments (BH-adjusted $p < 0.05$; top 3 per module, source, and phase) across alveolarization and mature-lung community gene sets.

| Phase | Module | Source | GO term | BH-adjusted $p$ |
|---|---|---|---|---|
| Alveolarization | 1 | GO-BP | innate immune response-activating signaling pathway | 9.24e-03 |
| Alveolarization | 1 | GO-BP | MyD88-dependent toll-like receptor signaling pathway | 9.24e-03 |
| Alveolarization | 1 | GO-BP | negative regulation of epidermis development | 9.24e-03 |
| Alveolarization | 1 | GO-CC | zymogen granule membrane | 9.36e-03 |
| Alveolarization | 1 | GO-MF | intracellularly calcium-gated channel activity | 1.04e-02 |
| Alveolarization | 1 | GO-MF | intracellularly calcium-gated chloride channel activity | 1.04e-02 |
| Alveolarization | 1 | GO-MF | oligosaccharide binding | 1.04e-02 |
| Alveolarization | 1 | GO-CC | microvillus | 1.88e-02 |
| Alveolarization | 1 | GO-CC | collagen-containing extracellular matrix | 4.00e-02 |

*Continued on next page*

*Table S5 continued*

| Phase | Module | Source | GO term | BH-adjusted $p$ |
|---|---|---|---|---|
| Alveolarization | 2 | GO-CC | MHC class II protein complex | 3.92e-10 |
| Alveolarization | 2 | GO-MF | MHC class II protein complex binding | 1.16e-09 |
| Alveolarization | 2 | GO-BP | antigen processing and presentation of exogenous peptide antigen via MHC class II | 1.31e-07 |
| Alveolarization | 2 | GO-BP | MHC class II protein complex assembly | 3.23e-07 |
| Alveolarization | 2 | GO-BP | peptide antigen assembly with MHC class II protein complex | 3.23e-07 |
| Alveolarization | 2 | GO-CC | collagen-containing extracellular matrix | 1.77e-06 |
| Alveolarization | 2 | GO-MF | extracellular matrix structural constituent | 2.08e-06 |
| Alveolarization | 2 | GO-MF | peptide antigen binding | 1.65e-04 |
| Alveolarization | 2 | GO-CC | late endosome membrane | 1.05e-03 |
| Alveolarization | 4 | GO-MF | methylated histone binding | 1.26e-04 |
| Alveolarization | 4 | GO-MF | histone binding | 3.20e-04 |
| Alveolarization | 5 | GO-MF | hydro-lyase activity | 1.25e-02 |
| Alveolarization | 5 | GO-MF | chitin binding | 2.68e-02 |
| Alveolarization | 5 | GO-MF | endopeptidase inhibitor activity | 2.68e-02 |
| Alveolarization | 5 | GO-CC | cytosolic ribosome | 3.80e-02 |
| Alveolarization | 5 | GO-CC | glycinergic synapse | 4.49e-02 |
| Alveolarization | 5 | GO-CC | peptidase inhibitor complex | 4.49e-02 |
| Alveolarization | 6 | GO-MF | ATP-dependent protein folding chaperone | 1.73e-02 |
| Alveolarization | 6 | GO-MF | fatty acid derivative binding | 1.73e-02 |
| Alveolarization | 6 | GO-MF | protein-containing complex destabilizing activity | 1.73e-02 |
| Alveolarization | 6 | GO-CC | endosome lumen | 2.37e-02 |
| Alveolarization | 6 | GO-CC | glycinergic synapse | 2.37e-02 |
| Alveolarization | 6 | GO-CC | lysosomal lumen | 2.37e-02 |
| Alveolarization | 6 | GO-BP | canonical inflammasome complex assembly | 2.47e-02 |
| Alveolarization | 6 | GO-BP | cardiac muscle cell apoptotic process | 2.47e-02 |
| Alveolarization | 6 | GO-BP | cell-cell fusion | 2.47e-02 |
| Mature Lung | 1 | GO-MF | estrogen 2-hydroxylase activity | 9.11e-04 |
| Mature Lung | 1 | GO-MF | aromatase activity | 4.95e-03 |
| Mature Lung | 1 | GO-MF | Hsp90 protein binding | 4.95e-03 |
| Mature Lung | 1 | GO-BP | long-chain fatty acid metabolic process | 1.05e-02 |
| Mature Lung | 1 | GO-BP | response to nutrient | 1.05e-02 |
| Mature Lung | 1 | GO-BP | xenobiotic catabolic process | 1.05e-02 |
| Mature Lung | 2 | GO-MF | methylated histone binding | 8.29e-03 |
| Mature Lung | 2 | GO-BP | humoral immune response | 1.28e-02 |
| Mature Lung | 2 | GO-CC | interstitial matrix | 1.53e-02 |
| Mature Lung | 2 | GO-CC | perisynaptic extracellular matrix | 1.53e-02 |
| Mature Lung | 2 | GO-CC | protein complex involved in cell-matrix adhesion | 1.53e-02 |
| Mature Lung | 2 | GO-MF | histone binding | 2.07e-02 |
| Mature Lung | 2 | GO-MF | calcium-dependent cysteine-type endopeptidase activity | 2.08e-02 |
| Mature Lung | 2 | GO-BP | disruption of cell in another organism | 2.40e-02 |
| Mature Lung | 2 | GO-BP | killing of cells of another organism | 2.40e-02 |
| Mature Lung | 3 | GO-CC | MHC class II protein complex | 2.26e-11 |
| Mature Lung | 3 | GO-MF | MHC class II protein complex binding | 1.01e-10 |
| Mature Lung | 3 | GO-BP | antigen processing and presentation of exogenous peptide antigen via MHC class II | 1.12e-08 |
| Mature Lung | 3 | GO-BP | MHC class II protein complex assembly | 2.99e-07 |
| Mature Lung | 3 | GO-BP | peptide antigen assembly with MHC class II protein complex | 2.99e-07 |
| Mature Lung | 3 | GO-CC | late endosome | 3.65e-06 |
| Mature Lung | 3 | GO-MF | peptide binding | 1.12e-05 |
| Mature Lung | 3 | GO-MF | CD4 receptor binding | 2.97e-05 |
| Mature Lung | 3 | GO-CC | lysosomal membrane | 1.53e-04 |

*Continued on next page*

*Table S5 continued*

| Phase | Module | Source | GO term | BH-adjusted $p$ |
|---|---|---|---|---|
| Mature Lung | 4 | GO-CC | cytosolic ribosome | 7.11e-05 |
| Mature Lung | 4 | GO-MF | structural constituent of ribosome | 9.06e-05 |
| Mature Lung | 4 | GO-BP | cytoplasmic translation | 8.03e-04 |
| Mature Lung | 4 | GO-BP | antimicrobial humoral immune response mediated by antimicrobial peptide | 1.34e-02 |
| Mature Lung | 4 | GO-BP | keratinocyte differentiation | 1.34e-02 |
| Mature Lung | 4 | GO-CC | peptidase inhibitor complex | 1.74e-02 |
| Mature Lung | 4 | GO-MF | oligosaccharide binding | 2.54e-02 |
| Mature Lung | 4 | GO-MF | peptidoglycan binding | 2.54e-02 |
| Mature Lung | 4 | GO-CC | A band | 2.66e-02 |
| Mature Lung | 5 | GO-CC | glycinergic synapse | 3.47e-03 |
| Mature Lung | 5 | GO-BP | endothelial cell proliferation | 1.01e-02 |
| Mature Lung | 5 | GO-BP | lymphocyte mediated immunity | 1.01e-02 |
| Mature Lung | 5 | GO-BP | negative regulation of toll-like receptor 4 signaling pathway | 1.01e-02 |
| Mature Lung | 5 | GO-MF | nuclear receptor activity | 2.19e-02 |
| Mature Lung | 5 | GO-CC | alpha-beta T cell receptor complex | 4.91e-02 |
| Mature Lung | 5 | GO-CC | dendritic spine | 4.91e-02 |
| Mature Lung | 6 | GO-MF | extracellular matrix structural constituent conferring compression resistance | 1.64e-04 |
| Mature Lung | 6 | GO-CC | collagen-containing extracellular matrix | 4.49e-03 |
| Mature Lung | 6 | GO-MF | carbonate dehydratase activity | 2.42e-02 |
| Mature Lung | 6 | GO-MF | chitin binding | 2.42e-02 |
| Mature Lung | 6 | GO-CC | alpha-beta T cell receptor complex | 2.74e-02 |
| Mature Lung | 6 | GO-CC | high-density lipoprotein particle | 2.92e-02 |

#### 2.7 Inference Argument Guidance

Table S6 summarizes practical recommendations for choosing PARROT scan and inference arguments depending on the analytical setting.

Table S6: Practical guidance for PARROT scan and inference arguments.

| Use case / condition | Suggested arguments | Strengths | Key assumptions and cautions |
|---|---|---|---|
| Highest localization accuracy (single CP; moderate $T$) | `scan_method="profile"` `compute_cis=TRUE` | Most faithful to the full profile objective; interpretable profile-based CI | Computationally expensive; each candidate split needs segment refits. CI uses a $\chi^2$ cutoff on the ELBO profile, so finite-sample coverage is approximate. |
| Long sequences / runtime-constrained screening | `scan_method="score"` | Substantially faster candidate scanning; practical for large $T$ | Score scan approximates profile refitting and may lose localization accuracy. If final localization matters, confirm with profile scan when feasible. |
| Robust inference under weak asymptotics | `pvalue_method=` `"permutation"` large `n_perm` | Fewer distributional assumptions than asymptotic $\chi^2$ calibration | Requires exchangeability of time labels under the null; temporal dependence can violate this. Resolution is limited by $1/(B+1)$ where $B$ is the permutation count (e.g., $B = 120 \Rightarrow p_{\min} = 0.00826$). |
| Model-based null calibration check | `pvalue_method=` `"bootstrap"` large `n_boot` | Adapts null calibration to fitted SBM parameter scale | Assumes the fitted null SBM is adequate; poor null fit can bias bootstrap calibration. High computational cost: each bootstrap replicate requires simulation and refitting. |
| Small sample / short segments | Increase `min_segment`; use fewer CPs; prefer profile + permutation | Stabilizes segment fits and reduces over-fragmentation | Very short segments can destabilize VEM, inflate uncertainty, and weaken p-value reliability. Report sensitivity analyses when segment lengths are close to the minimum. |

#### 2.8 Global, Community, and Mixed Change: Detailed Results

To clarify when PARROT excels, we simulated three controlled scenario families in the main benchmark: (i) global mean shift, (ii) pure community swap, and (iii) mixed transition (moderate global shift plus partial community rewiring). Detailed per-network tables are provided for all three families (Tables S8–S19), alongside an overall scenario-balanced ranking summary (Table S7).

##### 2.8.1 Failure modes and practical diagnostics

Performance is not uniform across regimes. Low-signal global Bernoulli settings (Tables S9 and S11) are difficult for all methods, including PARROT, and can yield weak detection and poor exact change-point localization; these are expected stress tests rather than contradictory evidence to rewiring-focused scenarios.

We provide an overall scenario-balanced ranking across scenario families and network types after the per-scenario breakdown; interpret this summary jointly with the per-scenario tables, not as a replacement.

Table S7: Overall method-balance summary across 3 scenarios, 4 network types, and 20 replicates per configuration. Lower Overall Rank indicates better aggregate performance (DetectionRate and F1 high, MAE low).

| Method | DetectionRate | F1 | ExactRate | MAE | Scenario Rank |
|---|---|---|---|---|---|
| PARROT | 0.742 | 0.742 | 0.596 | 1.412 | 1.25 |
| Subspace-CPD | 0.504 | 0.504 | 0.033 | 2.979 | 2.75 |
| ECP | 0.662 | 0.662 | 0.025 | 2.346 | 2.83 |
| Kernel-CPD | 0.242 | 0.242 | 0.242 | 3.792 | 4.00 |
| gSeg | 0.225 | 0.225 | 0.208 | 3.888 | 4.17 |

Table S8: Global Mean Shift: Unipartite Gaussian (N=12 nodes, T=10 time points, n=20 replicates).

| Method | Precision | Recall | F1 | MAE | Det. Rate | Exact |
|---|---|---|---|---|---|---|
| PARROT | $0.850 \pm 0.366$ | $0.850 \pm 0.366$ | $0.850 \pm 0.366$ | $0.80 \pm 1.82$ | 85.0% | 80.0% |
| Kernel-CPD | $0.000 \pm 0.000$ | $0.000 \pm 0.000$ | $0.000 \pm 0.000$ | $5.00 \pm 0.00$ | 0.0% | 0.0% |
| gSeg | $0.200 \pm 0.410$ | $0.200 \pm 0.410$ | $0.200 \pm 0.410$ | $4.00 \pm 2.05$ | 20.0% | 20.0% |
| ECP | $0.400 \pm 0.503$ | $0.400 \pm 0.503$ | $0.400 \pm 0.503$ | $3.40 \pm 2.01$ | 40.0% | 0.0% |
| Subspace-CPD | $0.850 \pm 0.366$ | $0.850 \pm 0.366$ | $0.850 \pm 0.366$ | $1.55 \pm 1.50$ | 85.0% | 5.0% |

Table S9: Global Mean Shift: Unipartite Bernoulli (N=12 nodes, T=10 time points, n=20 replicates).

| Method | Precision | Recall | F1 | MAE | Det. Rate | Exact |
|---|---|---|---|---|---|---|
| PARROT | $0.100 \pm 0.308$ | $0.100 \pm 0.308$ | $0.100 \pm 0.308$ | $4.50 \pm 1.28$ | 10.0% | 0.0% |
| Kernel-CPD | $0.000 \pm 0.000$ | $0.000 \pm 0.000$ | $0.000 \pm 0.000$ | $5.00 \pm 0.00$ | 0.0% | 0.0% |
| gSeg | $0.000 \pm 0.000$ | $0.000 \pm 0.000$ | $0.000 \pm 0.000$ | $4.90 \pm 0.45$ | 0.0% | 0.0% |
| ECP | $0.050 \pm 0.224$ | $0.050 \pm 0.224$ | $0.050 \pm 0.224$ | $4.75 \pm 1.12$ | 5.0% | 5.0% |
| Subspace-CPD | $0.150 \pm 0.366$ | $0.150 \pm 0.366$ | $0.150 \pm 0.366$ | $4.40 \pm 1.50$ | 15.0% | 5.0% |

Table S10: Global Mean Shift: Bipartite Gaussian (N=12 nodes, T=10 time points, n=20 replicates).

| Method | Precision | Recall | F1 | MAE | Det. Rate | Exact |
|---|---|---|---|---|---|---|
| PARROT | $0.350 \pm 0.489$ | $0.350 \pm 0.489$ | $0.350 \pm 0.489$ | $3.25 \pm 2.45$ | 35.0% | 35.0% |
| Kernel-CPD | $0.000 \pm 0.000$ | $0.000 \pm 0.000$ | $0.000 \pm 0.000$ | $5.00 \pm 0.00$ | 0.0% | 0.0% |
| gSeg | $0.200 \pm 0.410$ | $0.200 \pm 0.410$ | $0.200 \pm 0.410$ | $4.10 \pm 1.86$ | 20.0% | 10.0% |
| ECP | $0.450 \pm 0.510$ | $0.450 \pm 0.510$ | $0.450 \pm 0.510$ | $3.10 \pm 2.17$ | 45.0% | 10.0% |
| Subspace-CPD | $0.800 \pm 0.410$ | $0.800 \pm 0.410$ | $0.800 \pm 0.410$ | $1.75 \pm 1.68$ | 80.0% | 5.0% |

Table S11: Global Mean Shift: Bipartite Bernoulli (N=12 nodes, T=10 time points, n=20 replicates).

| Method | Precision | Recall | F1 | MAE | Det. Rate | Exact |
|---|---|---|---|---|---|---|
| PARROT | $0.250 \pm 0.444$ | $0.250 \pm 0.444$ | $0.250 \pm 0.444$ | $3.90 \pm 1.97$ | 25.0% | 10.0% |
| Kernel-CPD | $0.000 \pm 0.000$ | $0.000 \pm 0.000$ | $0.000 \pm 0.000$ | $5.00 \pm 0.00$ | 0.0% | 0.0% |
| gSeg | $0.050 \pm 0.224$ | $0.050 \pm 0.224$ | $0.050 \pm 0.224$ | $4.85 \pm 0.67$ | 5.0% | 0.0% |
| ECP | $0.150 \pm 0.366$ | $0.150 \pm 0.366$ | $0.150 \pm 0.366$ | $4.35 \pm 1.60$ | 15.0% | 5.0% |
| Subspace-CPD | $0.250 \pm 0.444$ | $0.250 \pm 0.444$ | $0.250 \pm 0.444$ | $4.05 \pm 1.73$ | 25.0% | 5.0% |

Table S12: Community Swap: Unipartite Gaussian (N=12 nodes, T=10 time points, n=20 replicates).

| Method | Precision | Recall | F1 | MAE | Det. Rate | Exact |
|---|---|---|---|---|---|---|
| PARROT | 0.900 ± 0.308 | 0.900 ± 0.308 | 0.900 ± 0.308 | 0.55 ± 1.05 | 90.0% | 75.0% |
| Kernel-CPD | 0.700 ± 0.470 | 0.700 ± 0.470 | 0.700 ± 0.470 | 1.50 ± 2.35 | 70.0% | 70.0% |
| gSeg | 0.350 ± 0.489 | 0.350 ± 0.489 | 0.350 ± 0.489 | 3.25 ± 2.45 | 35.0% | 35.0% |
| ECP | 1.000 ± 0.000 | 1.000 ± 0.000 | 1.000 ± 0.000 | 1.00 ± 0.00 | 100.0% | 0.0% |
| Subspace-CPD | 0.800 ± 0.410 | 0.800 ± 0.410 | 0.800 ± 0.410 | 1.85 ± 1.63 | 80.0% | 0.0% |

Table S13: Community Swap: Unipartite Bernoulli (N=12 nodes, T=10 time points, n=20 replicates).

| Method | Precision | Recall | F1 | MAE | Det. Rate | Exact |
|---|---|---|---|---|---|---|
| PARROT | 0.950 ± 0.224 | 0.950 ± 0.224 | 0.950 ± 0.224 | 0.50 ± 0.89 | 95.0% | 70.0% |
| Kernel-CPD | 0.000 ± 0.000 | 0.000 ± 0.000 | 0.000 ± 0.000 | 5.00 ± 0.00 | 0.0% | 0.0% |
| gSeg | 0.000 ± 0.000 | 0.000 ± 0.000 | 0.000 ± 0.000 | 5.00 ± 0.00 | 0.0% | 0.0% |
| ECP | 0.800 ± 0.410 | 0.800 ± 0.410 | 0.800 ± 0.410 | 1.80 ± 1.64 | 80.0% | 0.0% |
| Subspace-CPD | 0.150 ± 0.366 | 0.150 ± 0.366 | 0.150 ± 0.366 | 4.35 ± 1.60 | 15.0% | 5.0% |

Table S14: Community Swap: Bipartite Gaussian (N=12 nodes, T=10 time points, n=20 replicates).

| Method | Precision | Recall | F1 | MAE | Det. Rate | Exact |
|---|---|---|---|---|---|---|
| PARROT | 1.000 ± 0.000 | 1.000 ± 0.000 | 1.000 ± 0.000 | 0.20 ± 0.52 | 100.0% | 85.0% |
| Kernel-CPD | 0.550 ± 0.510 | 0.550 ± 0.510 | 0.550 ± 0.510 | 2.25 ± 2.55 | 55.0% | 55.0% |
| gSeg | 0.600 ± 0.503 | 0.600 ± 0.503 | 0.600 ± 0.503 | 2.00 ± 2.51 | 60.0% | 60.0% |
| ECP | 1.000 ± 0.000 | 1.000 ± 0.000 | 1.000 ± 0.000 | 1.00 ± 0.00 | 100.0% | 0.0% |
| Subspace-CPD | 0.850 ± 0.366 | 0.850 ± 0.366 | 0.850 ± 0.366 | 1.60 ± 1.50 | 85.0% | 5.0% |

Table S15: Community Swap: Bipartite Bernoulli (N=12 nodes, T=10 time points, n=20 replicates).

| Method | Precision | Recall | F1 | MAE | Det. Rate | Exact |
|---|---|---|---|---|---|---|
| PARROT | 0.900 ± 0.308 | 0.900 ± 0.308 | 0.900 ± 0.308 | 0.70 ± 1.30 | 90.0% | 65.0% |
| Kernel-CPD | 0.000 ± 0.000 | 0.000 ± 0.000 | 0.000 ± 0.000 | 5.00 ± 0.00 | 0.0% | 0.0% |
| gSeg | 0.050 ± 0.224 | 0.050 ± 0.224 | 0.050 ± 0.224 | 4.75 ± 1.12 | 5.0% | 5.0% |
| ECP | 0.800 ± 0.410 | 0.800 ± 0.410 | 0.800 ± 0.410 | 1.80 ± 1.67 | 80.0% | 5.0% |
| Subspace-CPD | 0.100 ± 0.308 | 0.100 ± 0.308 | 0.100 ± 0.308 | 4.55 ± 1.39 | 10.0% | 5.0% |

Table S16: Mixed Transition: Unipartite Gaussian (N=12 nodes, T=10 time points, n=20 replicates).

| Method | Precision | Recall | F1 | MAE | Det. Rate | Exact |
|---|---|---|---|---|---|---|
| PARROT | 1.000 ± 0.000 | 1.000 ± 0.000 | 1.000 ± 0.000 | 0.00 ± 0.00 | 100.0% | 100.0% |
| Kernel-CPD | 0.800 ± 0.410 | 0.800 ± 0.410 | 0.800 ± 0.410 | 1.00 ± 2.05 | 80.0% | 80.0% |
| gSeg | 0.450 ± 0.510 | 0.450 ± 0.510 | 0.450 ± 0.510 | 2.75 ± 2.55 | 45.0% | 45.0% |
| ECP | 1.000 ± 0.000 | 1.000 ± 0.000 | 1.000 ± 0.000 | 1.00 ± 0.00 | 100.0% | 0.0% |
| Subspace-CPD | 0.900 ± 0.308 | 0.900 ± 0.308 | 0.900 ± 0.308 | 1.50 ± 1.24 | 90.0% | 0.0% |

Table S17: Mixed Transition: Unipartite Bernoulli (N=12 nodes, T=10 time points, n=20 replicates).

| Method | Precision | Recall | F1 | MAE | Det. Rate | Exact |
|---|---|---|---|---|---|---|
| PARROT | 0.700 ± 0.470 | 0.700 ± 0.470 | 0.700 ± 0.470 | 1.80 ± 2.04 | 70.0% | 35.0% |
| Kernel-CPD | 0.000 ± 0.000 | 0.000 ± 0.000 | 0.000 ± 0.000 | 5.00 ± 0.00 | 0.0% | 0.0% |
| gSeg | 0.050 ± 0.224 | 0.050 ± 0.224 | 0.050 ± 0.224 | 4.75 ± 1.12 | 5.0% | 5.0% |
| ECP | 0.500 ± 0.513 | 0.500 ± 0.513 | 0.500 ± 0.513 | 3.15 ± 1.93 | 50.0% | 0.0% |
| Subspace-CPD | 0.050 ± 0.224 | 0.050 ± 0.224 | 0.050 ± 0.224 | 4.85 ± 0.67 | 5.0% | 0.0% |

Table S18: Mixed Transition: Bipartite Gaussian (N=12 nodes, T=10 time points, n=20 replicates).

| Method | Precision | Recall | F1 | MAE | Det. Rate | Exact |
|---|---|---|---|---|---|---|
| PARROT | 0.900 ± 0.308 | 0.900 ± 0.308 | 0.900 ± 0.308 | 0.50 ± 1.28 | 90.0% | 80.0% |
| Kernel-CPD | 0.850 ± 0.366 | 0.850 ± 0.366 | 0.850 ± 0.366 | 0.75 ± 1.83 | 85.0% | 85.0% |
| gSeg | 0.600 ± 0.503 | 0.600 ± 0.503 | 0.600 ± 0.503 | 2.00 ± 2.51 | 60.0% | 60.0% |
| ECP | 1.000 ± 0.000 | 1.000 ± 0.000 | 1.000 ± 0.000 | 1.00 ± 0.00 | 100.0% | 0.0% |
| Subspace-CPD | 1.000 ± 0.000 | 1.000 ± 0.000 | 1.000 ± 0.000 | 1.05 ± 0.22 | 100.0% | 0.0% |

Table S19: Mixed Transition: Bipartite Bernoulli (N=12 nodes, T=10 time points, n=20 replicates).

| Method | Precision | Recall | F1 | MAE | Det. Rate | Exact |
|---|---|---|---|---|---|---|
| PARROT | 1.000 ± 0.000 | 1.000 ± 0.000 | 1.000 ± 0.000 | 0.25 ± 0.55 | 100.0% | 80.0% |
| Kernel-CPD | 0.000 ± 0.000 | 0.000 ± 0.000 | 0.000 ± 0.000 | 5.00 ± 0.00 | 0.0% | 0.0% |
| gSeg | 0.150 ± 0.366 | 0.150 ± 0.366 | 0.150 ± 0.366 | 4.30 ± 1.72 | 15.0% | 10.0% |
| ECP | 0.800 ± 0.410 | 0.800 ± 0.410 | 0.800 ± 0.410 | 1.80 ± 1.67 | 80.0% | 5.0% |
| Subspace-CPD | 0.150 ± 0.366 | 0.150 ± 0.366 | 0.150 ± 0.366 | 4.25 ± 1.62 | 15.0% | 5.0% |

#### 3 Software and Data Availability

All GEO accessions were retrieved from NCBI GEO. Plotting used `ggplot2` and alluvial visualizations used `ggalluvial`; GO enrichment annotations used `clusterProfiler`.

##### 3.1 PARROT R Package

PARROT (version 1.0.0) is available as an R package. Key exported functions:

- `parrot()`: End-to-end change point detection with automatic network-type inference, ICL-based $Q$ selection, and optional p-values/confidence intervals
- `simulate_sbm()`, `simulate_sbm_cp()`, `simulate_sbm_multi_cp()`: Simulate static, single-CP, or multi-CP network sequences (unipartite/bipartite $\times$ Gaussian/Bernoulli)
- `fit_sbm()`: Variational EM estimation of SBM parameters with ICL model selection
- `detect_single_cp()`,`detect_multiple_cp()`: Score-scan or profile-likelihood detection with WBS/BS support
- `compute_pvalue()`: P-value computation via Wilks, permutation, or bootstrap tests
- `plot.parrot_result()`,`plot_network_heatmap()`,`plot_network_evolution()`: Visualization of results and network dynamics

##### 3.2 Installation

```
# Install from source
R CMD INSTALL PARROT_1.0.0.tar.gz

# Or from GitHub
devtools::install_github("cchen22/PARROT")
```

##### 3.3 Example Usage

```
library(PARROT)

# Simulate data
data <- simulate_sbm_cp(
  N = 40, T_len = 60, cp_time = 30, Q = 2,
  type = "unipartite", distribution = "gaussian",
  theta1 = list(mu = matrix(c(2, 0, 0, 2), 2, 2), sigma = 0.3),
  theta2 = list(mu = matrix(c(0, 2, 2, 0), 2, 2), sigma = 0.3)
)

# Detect change points
result <- parrot(data$Y, Q = 2, distribution = "gaussian")
print(result)
plot(result)
```

##### 3.4 Reproducibility

All manuscript figure-generation scripts are available:

- `reproduce_all_outputs.R`: one-command entrypoint to regenerate all main/supplementary figure and table outputs
- `generate_figure1_overview.R`: Figure 1 (method overview and pipeline illustration)
- `reproduce_simulation_global_vs_community.R`: simulation benchmark generation and Supplementary Tables S7–S19
- `generate_figure2.R`: Figure 2 multi-panel visualization from `Figure2_AllResults.csv`
- `generate_figS1_sbm_recovery.R`: Supplementary Figure S1 (SBM parameter recovery)
- `generate_figS2_q_selection_realdata.R`: Supplementary Figure S2 (real-data $Q$ selection via ICL)
- `data_utils_realdata.R`: shared loaders/caching for preprocessed GEO real-data analyses
- `generate_figure3_cardiac_scrna.R`: Figure 3 (human hiPSC cardiac differentiation real-data application; GSE202398 scRNA-seq)
- `generate_figure4_lung_development.R`: Figure 4 (mouse postnatal lung development real-data application; GSE74243)
- `generate_figS3_score_vs_profile_scan.R`: Supplementary Figure S3 (score-scan vs profile-scan benchmark)
- `generate_tableS3_cardiac_robustness.R`: Supplementary Table S3 (cardiac sensitivity checks)
- `generate_tableS4_lung_robustness.R`: Supplementary Table S4 (lung sensitivity checks)
- `generate_tableS5_go_enrichment_lung.R`: Supplementary Table S5 (lung GO enrichment summary)
